# *Lin28a/let-7* Pathway Modulates the *Hox* Code via *Polycomb* Regulation during Axial Patterning in Vertebrates

**DOI:** 10.1101/854091

**Authors:** Tempei Sato, Kensuke Kataoka, Yoshiaki Ito, Shigetoshi Yokoyama, Masafumi Inui, Masaki Mori, Satoru Takahashi, Keiichi Akita, Shuji Takada, Hiroe Ueno-Kudoh, Hiroshi Asahara

**Affiliations:** Department of Systems BioMedicine, Graduate School of Medical and Dental Sciences, Tokyo Medical and Dental University, Tokyo 113-8510, Japan; Department of Systems BioMedicine, National Research Institute for Child Health and Development, Tokyo 157-8535, Japan; Research Fellow of Japan Society for the Promotion of Science, Tokyo, Japan; Research Core, Tokyo Medical and Dental University, Tokyo, 113-8510, Japan; Laboratory of Metabolism, National Institutes of Health, Bethesda, MD 20892-4255, USA; Laboratory of Animal Regeneration Systemology, Meiji University, Kanagawa, 214-8571, Japan; Department of Medical Chemistry, Shiga University of Medical Science, Shiga, 520-2192, Japan; Department of Anatomy and Embryology, University of Tsukuba, Ibaraki, 305-8575, Japan; Department of Clinical Anatomy, Graduate School of Medical and Dental Sciences, Tokyo Medical and Dental University, Tokyo 113-8510, Japan; Reproduction Center, Yokohama City University, Yokohama, 232-0024, Japan; AMED-CREST, Japan Agency for Medical Research and Development (AMED), 1-7-1 Otemachi, Chiyoda-ku, Tokyo 100-0004, Japan; Department of Molecular Medicine, The Scripps Research Institute, 10550 N. Torrey Pines Rd., La Jolla, CA 92037, USA

## Abstract

The body plan along the anteroposterior axis and regional identities are specified by the spatiotemporal expression of *Hox* genes. Multistep controls are required for their unique expression patterns; however, the molecular mechanisms behind the tight control of *Hox* genes are not fully understood. In this study, we demonstrated that the *Lin28a*/*let-7* reciprocal regulatory pathway is critical for vertebral specification. *Lin28a*^−/−^ mice exhibited homeotic transformations of vertebrae which were caused by the global dysregulation of posterior *Hox* genes. The accumulation of *let-7*-family microRNAs in *Lin28a*^−/−^ mice resulted in the reduction of PRC1 occupancy at the *Hox* cluster loci by targeting *Cbx2*. Consistently, Lin28a loss in embryonic stem-like cells led to aberrant induction of posterior *Hox* genes, which was rescued by the knockdown of *let-7*-family microRNAs. These results suggest that *Lin28*/*let-7* pathway is possibly involved in the modulation of the “*Hox* code” via *Polycomb* regulation during axial patterning in vertebrates.

## Introduction

The precise positioning of each organ and tissue has to be tightly controlled during embryogenesis. The body plan along the anteroposterior axis is modulated by the spatiotemporal expression of *Hox* genes, which is known as the “*Hox* code” (Wellik, 2007, Mallo and Alonso, 2013). *Hox* gene mutants show homeotic transformations in which a specific vertebra is morphologically changed into an anterior one, suggesting that the spatiotemporal expression patterns of *Hox* genes specify regional anatomical identities (Wellik, 2007). Multistage controls, such as transcriptional, posttranscriptional, and epigenetic regulation, are required for the nested expression patterns of *Hox* genes (Mallo and Alonso, 2013).

As for epigenetic control, *Polycomb* group (PcG) genes are involved in *Hox* gene regulation via the chromatin architecture at *Hox* cluster loci in a developmental time-dependent manner (Soshnikova, 2014). PcG genes form two complexes, the *Polycomb* Repressive Complex (PRC) 1 and PRC2. PRC2 includes Ezh2 which can catalyze H3K27me3 at target loci, and consequently, this specific histone modification causes the recruitment of PRC1 via Cbx2 in the complex to silence gene expression. Thus, the accumulation of PcG complexes at *Hox* clusters during embryogenesis leads to the transcriptional silencing of *Hox* genes, which is supported by evidence that the ablation of PcG genes dysregulates *Hox* gene expression, resulting in subsequent homeotic transformation in anteroposterior patterning (Mallo and Alonso, 2013, Soshnikova, 2014). During embryogenesis, PcG gene expression gradually diminished (Hashimoto et al., 1998), which leads to the initiation of spatiotemporal *Hox* gene expression. However, the molecular mechanisms underlying the termination of PcG gene expression remain largely unclear.

Previously, we generated a whole-mount *in situ* hybridization database called “EMBRYS” that covers ∼1,600 transcription factors and RNA-binding factors using mice at embryonic day (E)9.5, E10.5, and E11.5 (Yokoyama et al., 2009). Among these data, we were particularly interested in dynamic expressional changes of *Lin28a* during embryogenesis: at E9.5, *Lin28a* is expressed ubiquitously, whereas its expression gradually diminishes from head to tail at E10.5 and E11.5 (Yokoyama et al., 2008, Yokoyama et al., 2009). These unique expressional changes prompted us to analyze if *Lin28a* is involved in the spatiotemporal regulation of *Hox* genes.

*Lin-28* was identified as a heterochronic gene that regulates the developmental timing of multiple organs in *Caenorhabditis elegans* (*C.elegans*) (Moss et al., 1997). *Lin-28* encodes an RNA-binding protein, and the loss of function of *Lin-28* causes precocious development, with skipping of events that are specific to the second larval stage (Ambros and Horvitz, 1984, Moss et al., 1997). In contrast, mutants of *let-7*, a microRNA-encoding heterochronic gene, exhibit reiteration of the fourth larval developmental stage because of failures in terminal differentiation and cell-cycle exit (Pasquinelli et al., 2000, Reinhart et al., 2000). Importantly, *Lin-28* and *let-7* form a negative feedback loop that is essential for developmental timing in *C. elegans*. This reciprocal regulation between Lin28a and *let-7* is well conserved in mammals (Moss and Tang, 2003, Viswanathan et al., 2008); Lin28a promotes the degradation of *let-7* precursors (Heo et al., 2009, Chang et al., 2013), whereas *let-7* inhibits *Lin28a* expression via posttranscriptional regulation (Moss and Tang, 2003).

Vertebrates possess two homologous of *Lin28* genes, *Lin28a* and *Lin28b*. *Lin28a* is highly expressed in pluripotent stem cells and is ubiquitously expressed in the early embryonic stage, and its expression is diminished during development (Yang and Moss, 2003, Shyh-Chang and Daley, 2013, Yokoyama et al., 2008, Yokoyama et al., 2009). In contrast, *Lin28b* is dominantly expressed in testes, placenta, and fetal liver, as well as in undifferentiated hepatocarcinoma (Guo et al., 2006). The versatile functions of *Lin28a* are observed in diverse events such as germ layer formation (Faas et al., 2013), germ cell development (West et al., 2009), neural development (Yang et al., 2015), and glucose metabolism (Zhu et al., 2011). Conversely, *let-7*-family genes are highly expressed in differentiated tissues, and their products function as tumor suppressors via the inhibition of oncogenes such as *c-Myc*, *K-ras*, and *Hmga2* (Mayr et al., 2007, Lee and Dutta, 2007, Johnson et al., 2005, Sampson et al., 2007). These observations prompted us to test the potential “heterochronic” function of *Lin28a* in vertebrate development; however, it remains largely unclear if the evolutionally fundamental function of the *Lin-28* and *let-7* negative feedback loop in the regulation of developmental timing and pattern of *C. elegans* is conserved or adapted to in vertebrates.

In this work, we originally generated *Lin28a* knockout (*Lin28a*^−/−^) mice and analyzed the function of this gene in developmental patterning. We showed that *Lin28a*/*let-7* pathway is critical for axial patterning and vertebral specification. *Lin28a*^−/−^ mice exhibited homeotic transformations of vertebrae, which were accompanied by the global dysregulation of posterior *Hox* genes. The accumulation of *let-7*-family microRNAs in *Lin28a*^−/−^ mice resulted in the reduction of PRC1 occupancy at the *Hox* cluster loci by targeting *Cbx2*. Consistent with these results, Lin28a loss in embryonic stem-like cells led to the aberrant induction of posterior *Hox* genes, which was rescued by knockdown of *let-7*-family microRNAs. These results suggest the involvement of *Lin28*/*let-7* pathway in the modulation of the “*Hox* code” in vertebrates.

## Results

### *Lin28a*^−/−^ mice exhibit homeotic transformations

*Lin28a* exhibits unique spatiotemporal expression changes during early development (Figure 1A) (Yang and Moss, 2003; Yokoyama et al., 2008, 2009). At E9.5, *Lin28a* is expressed ubiquitously; and subsequently, its expression disappears from head to tail at around E10.5 and E11.5 (Yokoyama et al., 2008, 2009). To examine the potential significance of these dynamic expressional changes and of the developmental function of *Lin28a* in mice, we generated *Lin28a*^−/−^ mice (Figure S1). The normal Mendelian ratio of genotypes was observed for *Lin28a*^−/−^ mice during early to middle embryogenesis. However, the frequency of *Lin28a*^−/−^ mice decreased from E17.5 and after birth. Most of the *Lin28a*^−/−^ mice died perinatally or within a few days after birth (Table S1). *Lin28a*^−/−^ mice exhibited short stature compared with wild-type (Wt) mice and showed severe growth defects (Figure S2). These findings are consistent with previous reports that *Lin28a* is necessary for normal growth (Shinoda et al., 2013); however, our *Lin28a*^−/−^ mice showed severe phenotypes that might have been caused by differences in gene targeting construct and genetic background.

**Figure 1.**
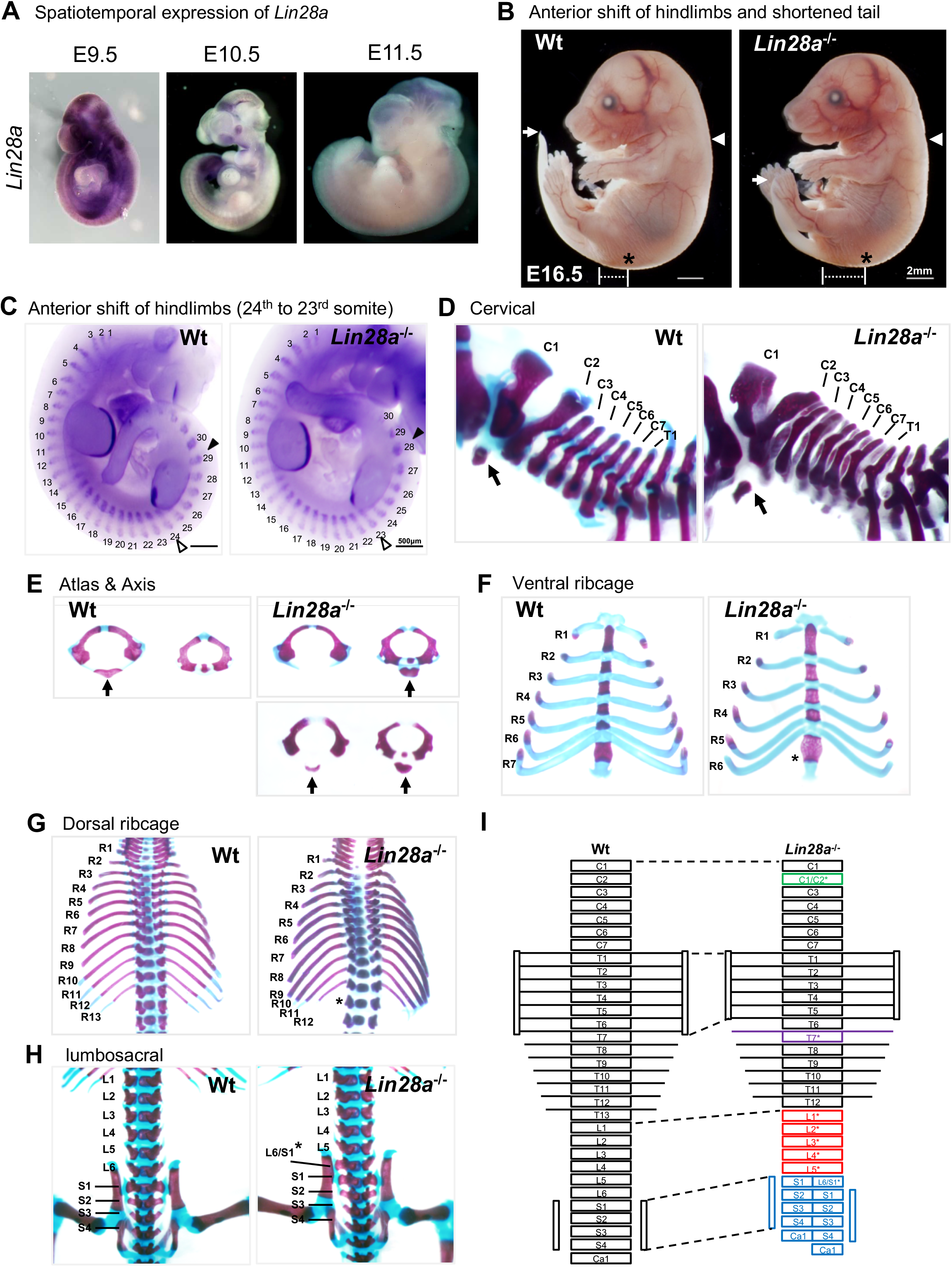
Homeotic transformations of vertebrae in *Lin28a*^-/-^ mice. (**A**) Whole-mount *in situ* hybridization of *Lin28a* in E9.5-11.5 embryos. (**B**) Lateral views of Wt (left panel) and *Lin28a*^−/−^ mice (right panel) at E16.5. White arrow, the tip of the tail; white arrowhead, forelimb position; asterisk, hindlimb position. (**C**) Whole-mount *in situ* hybridization of *Myog* and *FGF8* in E10.5 embryos. The numbers indicate the expression domains of *Myog*. White arrowhead, the starting position of the hindlimb bud; black arrowhead, the ending position of the hindlimb bud. (**D–H**) Representative skeletal preparations of Wt (left panels) and *Lin28a*^−/−^ mice (right panels). Lateral views of cervical and upper thoracic vertebrae (D); anterior views of the atlas and the axis (E); ventral views of the ribcage (F); dorsal views of thoracic vertebrae and ribs (G); and dorsal views of lumbar and sacral vertebrae (H) are shown. C1–C7, 1^st^ to 7^th^ cervical vertebrae; T1 and T2, 1^st^ and 2^nd^ thoracic vertebrae; R1–R13, 1^st^ to13^th^ ribs; L1–L6, 1^st^ to 6^th^ lumbar vertebrae; S1–S4, 1^st^ to 4^th^ sacral vertebrae; black arrows, anterior arch of the atlas. (**I**) schematic diagram of skeletal phenotypes in *Lin28a*^-/-^ mice.

We then examined anteroposterior axis formation in *Lin28a*^−/−^ mice since *Lin28a*^−/−^ mice showed a slight anterior shift of the hindlimbs and shortened tails (Figure 1B). To define the details of these phenotypes, whole-mount *in situ* hybridization of *Myog* and *Fgf8* was performed to outline somites and limb buds. The hindlimbs of *Lin28a*^−/−^ mice shifted anteriorly by one somite (from the 23^rd^ to the 28^th^ expression domains of *Myog*), whereas the position of the forelimb buds of *Lin28a*^−/−^ mice were not altered (Figure 1C). To analyze the potential functions of *Lin28a* in skeletal patterning, Alcian blue and Alizarin red S staining were applied to the skeletal preparations. Although bone and cartilage development was normal, homeotic transformations of vertebrae were observed in *Lin28a*^−/−^ mice (Figures 1D–1H). In *Lin28a*^−/−^ mice, the anterior arch of the atlas was formed from the second cervical vertebra (C2) (Figure 1D), and not from C1, as normally observed, or from the fusion of C1 and C2 (Figure 1E). These transformations were observed in 64.3 % of *Lin28a*^−/−^ mice and in 21.1 % of *Lin28a*^+/–^ mice; in contrast, they were never found in Wt mice (Table 1). There were only six pairs of true ribs attached to the sternum in *Lin28a*^−/−^ mice, whereas Wt and *Lin28a*^+/–^ mice had seven pairs (Figure 1F). Furthermore, an abnormal number of ribs was observed in *Lin28a*^−/−^ mice at 100 % penetrance, whereas Wt and *Lin28a*^+/–^ mice exhibited the normal 13 pairs of ribs (Figure 1G and Table 1). These results suggest that posterior transformations of vertebral identity occur in the 7^th^ and 13^th^ thoracic vertebrae during skeletal patterning. Moreover, partial transformations were observed in the first sacral vertebra (S1), producing a morphological feature of lumbar vertebrae on only one side (Figure 1H). The frequency of these observations was significantly higher in *Lin28a*^−/−^ mice (Table 1). Finally, *Lin28a*^−/−^ mice showed various homeotic transformations (Figure 1I), suggesting that *Lin28a* plays a critical role in the specification of vertebrae along the anteroposterior axis.

**Table 1.** Summary of skeletal abnormalities in *Lin28a* mutant mice.

|  | Anterior arch<br>of the atlas<sup>a</sup> | Ribs<sup>b</sup> |  | Sternum attachment<sup>c</sup> |  | Lumbar<sup>d</sup> |  |  |
| --- | --- | --- | --- | --- | --- | --- | --- | --- |
|  |  | 13 | 12 | 7 | 6 | 6 | 5 | L6/S1* |
| Wt (n = 16) | 0 | 16 (100%) | 0 | 16 (100%) | 0 | 7 (43.8%) | 8 (50%) | 1 (6.2%) |
| *Lin28a*<sup>+/−</sup><br>(n = 19) | 4 (21.1%) | 19 (100%) | 0 | 19 (100%) | 0 | 0 | 18 (94.7%) | 1 (5.3%) |
| *Lin28a*<sup>−/−</sup><br>(n = 14) | 9 (64.3%) | 0 | 14 (100%) | 0 | 14 (100%) | 0 | 9 (64.3%) | 5 (35.7%) |

The percentages of each phenotype are shown in parenthesis.
<sup>a</sup> The anterior arch of the atlas was formed from C2 or via fusion.
<sup>b</sup> Total number of pairs of ribs.
<sup>c</sup> Total number of pairs of true ribs that were attached to the sternum.
<sup>d</sup> Total number of lumbar vertebrae. L6/S1\* indicates an abnormal sacral vertebra that had morphological features of a lumbar vertebra on only one side.

### *Hox* genes are dysregulated in *Lin28a*^−/−^ mice

The morphologies and characteristics of each vertebra are specified by the spatiotemporal expression of *Hox* genes (Wellik, 2007). It was remarkable that *Lin28a*^−/−^ mice exhibited global transformations with high penetration, whereas mutants of *Hox* genes showed abnormalities in a limited region of vertebrae. To test if *Hox* genes are involved in the phenotypes found in *Lin28a*^−/−^ mice, we examined *Hox* gene expression during embryogenesis. Quantitative real-time polymerase chain reaction (q-PCR) analyses of *Hox* genes were performed at E9.5, a time at which *Lin28a* was ubiquitously expressed in Wt mice (Figure 1A). *Lin28a*^−/−^ mice exhibited global dysregulation of *Hox* genes, which was most remarkable for the 5′ (posterior) *Hox* genes (Figure 2A). Whole-mount *in situ* hybridization analyses revealed that the expression domain of *Hoxc13* and *Hoxd12* was enlarged anteriorly (Figures 2B, 2C, and S3). In contrast, there were no significant changes in the expression domain of the the other *Hox* genes upregulated in *Lin28a*^−/−^ mice (Hoxa3, d3, b8, c8, a11, and a13) (Figure S3). These results suggest that the homeotic transformations of vertebrae observed in *Lin28a*^−/−^ mice were caused by the upregulation of posterior *Hox* genes and/or the anteriorization of *Hox* gene expression.

**Figure 2.**
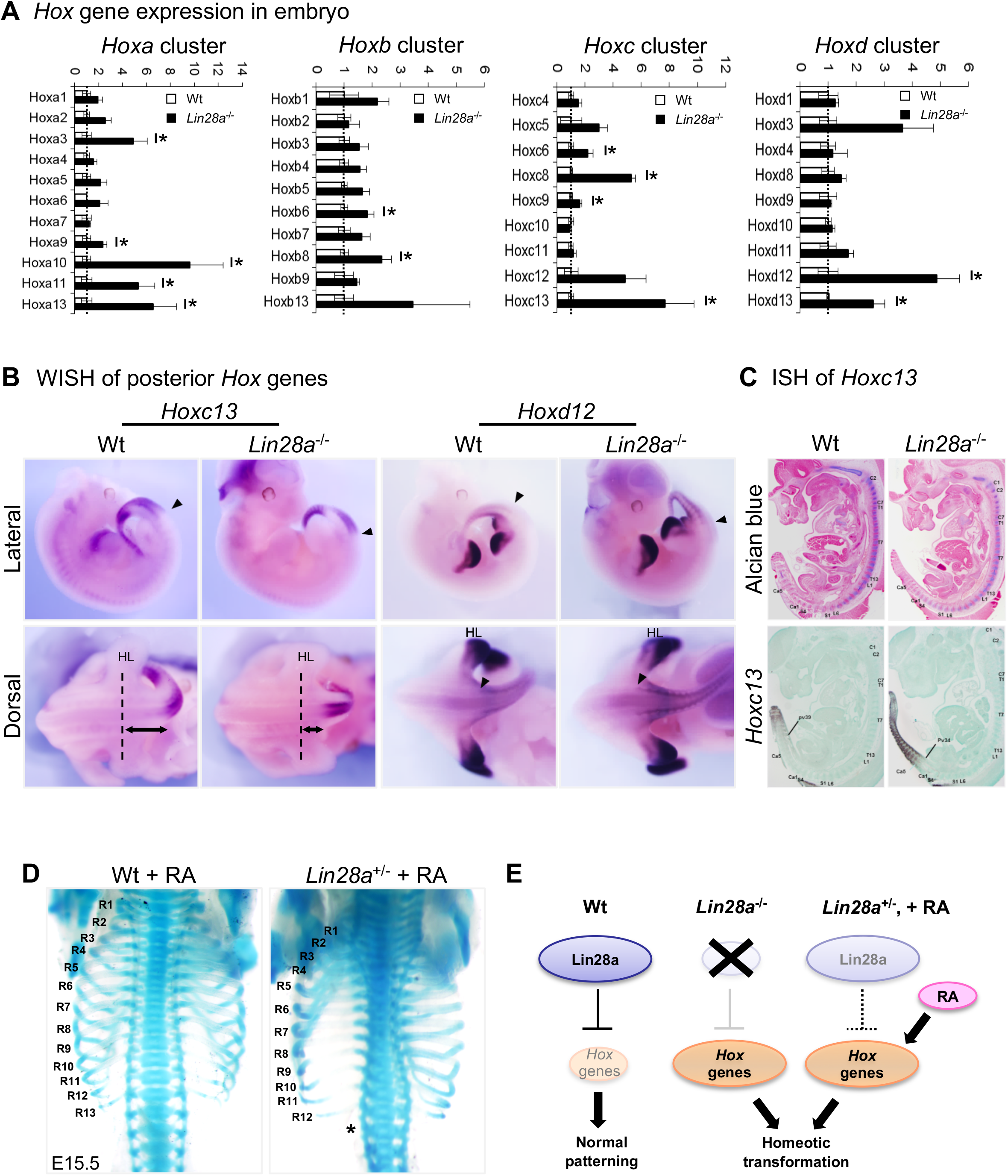
*Hox* gene dysregulation in *Lin28a*^-/-^ mice. (**A**) q-PCR analyses of all *Hox* genes. All data are expressed as the mean ± standard error of the mean (SEM) (n = 3). \**p* < 0.05. (**B**) Whole-mount *in situ* hybridization of *Hox* genes in E11.5 embryos. Lateral views (top panels) and dorsal views (bottom panels) of hindlimb and tail region are shown. Black arrowhead, anterior domain of *Hox* gene; HL, hindlimb; dashed line, hindlimb position; two-way arrow, distance from the hindlimb to the anterior domain of *Hoxc13*. (**C**) Histological analysis of E12.5 animals. Alcian blue staining (top panels) and *in situ* hybridization of *Hoxc13* (bottom panels) are shown. (**D**) Skeletal preparations of Wt (left panel) and *Lin28a*^+/–^ mice (right panel) that received RA treatment. R1–R13, 1^st^ to 13^th^ ribs; asterisk, the ablation of the 13^th^ rib. See also Figure S4. (**E**) Summary of *Hox* gene dysregulation in *Lin28a* mutants.

**Figure 3.**
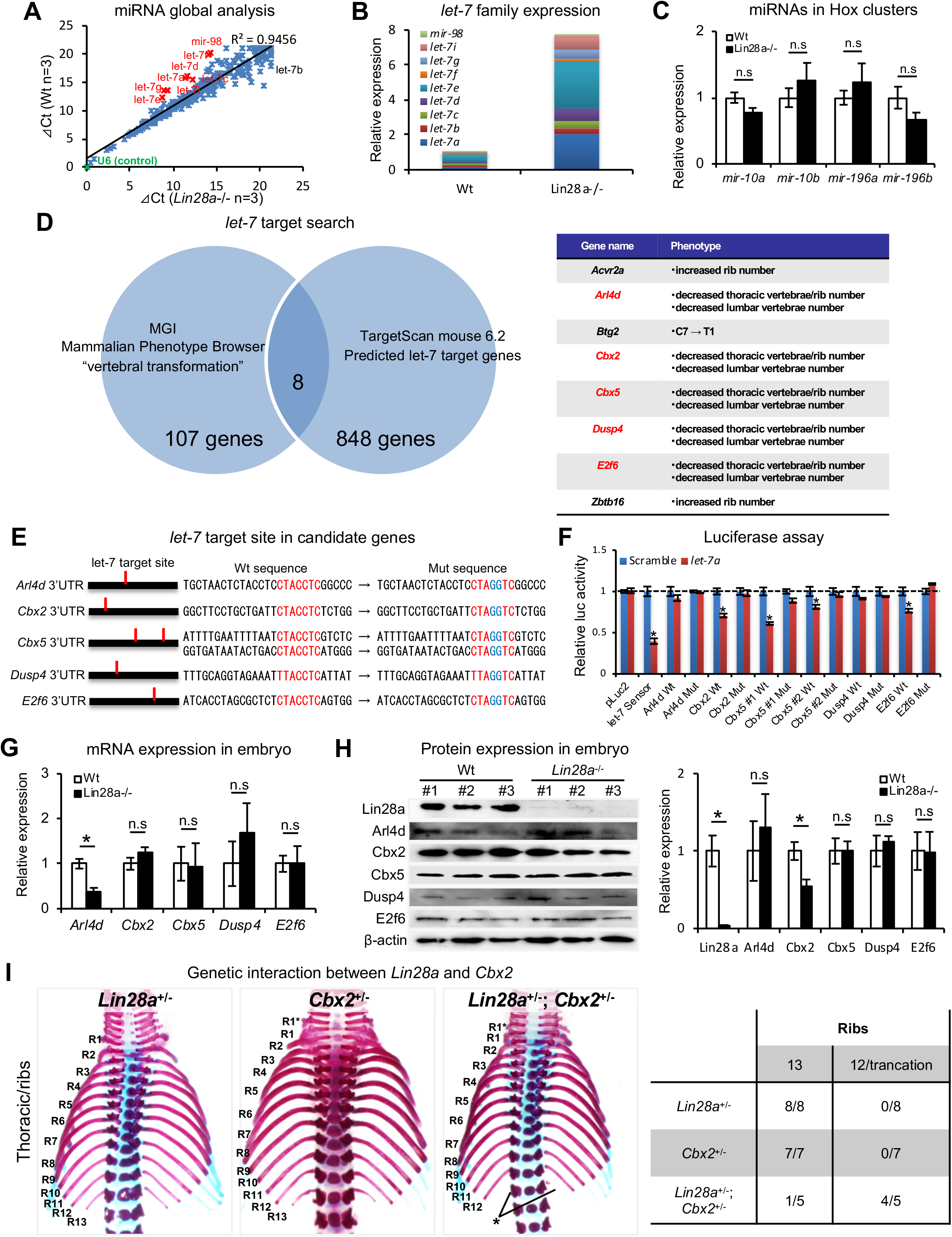
*Let-7* targets the polycomb gene directly. (**A**) Comparison of microRNA expression in Wt and *Lin28a*^−/−^ mice. (**B, C**) q-PCR analyses of *let-7*-family members (B) and *Hox*-embedded microRNAs (C). In (B), data are expressed as the mean (n = 3), and the relative amount of total *let-7* microRNAs is shown. (**D**) *let-7* target search with TargetScan and Phenotype Browser. (**E**) The *let-7* target site in the 3’UTR sequence of candidate genes. The *let-7* seed-matched sequence and mutated sequence are shown in red and blue, respectively. (**F**) Luciferase reporter activity in the presence/absence of the *let-7* target site in 3’UTR sequence. (**G-H**) qPCR and western blot analyses of candidate genes. (**I**) Dorsal views of thoracic vertebrae and ribs. Single heterozygous mutants (left and middle panels) and a double heterozygous mutant (right panel) are shown. R1–R13, 1^st^ to 13th ribs; asterisk, the ablation or truncation of the 13^th^ rib. See also Figure S5. (**J**) Frequency of rib defects in mutant mice. All data are expressed as the mean ± SEM (n = 3). * *p* < 0.05. n.s., not significant.

It is known that Hox genes are modulated by retinoic acid (RA) signaling, and that RA exposure causes posterior transformations of vertebrae via global anteriorization of *Hox* gene expression (Kessel and Gruss, 1991). Therefore we administered RA to pregnant *Lin28a*^+/–^ animals to test if this would augment the skeletal phenotypes observed in *Lin28a* mutants. RA was injected intraperitoneally at 7.5 days postcoitum (dpc) and the skeletal patterning of each fetus was analyzed. We found that *Lin28a* mutant embryos showed RA sensitivity. *Lin28a*^+/–^ mice that received RA treatment showed loss of the 13^th^ pair of ribs, which coincided with the findings observed in *Lin28a*^−/−^ mice, whereas no obvious defects were observed in Wt littermates (Figure 2D). In contrast, no additional defects in the thoracic region were observed in the *Lin28a*^−/−^ embryos that natively had only 12 pairs of ribs (Figure S4A). In the cervical region, the severity of skeletal patterning defects correlated with the genotype of *Lin28a* (Figures S4B–S4F). After RA treatment, some *Lin28a*^+/–^ embryos exhibited the C1/C2 fusion phenotype (Figure S4D), whereas *Lin28a*^−/−^ embryos showed more severe defects that were characterized by fusion of the exoccipital bone with C1 and C2 (Figures S4E and S4F). Taken together, these observations indicate that *Lin28a* acts upstream of the *Hox* genes, and that dysregulation of *Hox* genes might be responsible for the phenotype of *Lin28a*^−/−^ mice (Figure 2E).

### *Lin28a* regulates *Cbx2* expression via *let-7* repression

We examined the molecular mechanism underlying the *Lin28a-*mediated regulation of *Hox* gene expression during embryogenesis. Since *Lin28a* is known as a negative regulator of *let-7* biogenesis via TUT4-mediated terminal uridylation and inhibition of Dicer processing (Heo et al., 2009, Chang et al., 2013), we examined the microRNA expression profile of *Lin28a*^−/−^ mice. TaqMan microRNA array analyses were performed on E9.5 embryos. Consistent with previous reports (Viswanathan et al., 2008, Rybak et al., 2008), we found that mature microRNAs of *let-7*-family members were significantly accumulated in *Lin28a*^−/−^ mice (Figure 3A). These results were also confirmed by q-PCR analysis of *let-7*-family members (Figure 3B). There was no difference between Wt and *Lin28a*^−/−^ mice with regards to the expression of *mir-10s* and *mir-196s*, which are regulators of the spatial expression of *Hox* genes and of vertebral specification (Woltering and Durston, 2008, Yekta et al., 2004, Hornstein et al., 2005) (Figure 3C). Consistent with previous reports (Heo et al., 2009, Chang et al., 2013), these results imply that the ablation of Lin28a promotes the specific accumulation of *let-7* family microRNAs during embryogenesis.

We then sought a potential target gene for *let-7*, which might have been involved in the homeotic transformations observed in *Lin28a*^−/−^ mice. We performed comprehensive screening for the *let-7* target candidate genes using the following criteria; 1) *let-7* target genes, computationally predicted using TargetScan (856 genes) and 2) Annotated genes responsible for posterior transformations and of which knockout mice show vertebrae that are similar to those observed in *Lin28a*^-/-^ mice, as screened by Mouse Genome Informatics (115 genes). We found that five of the genes (*Arl4d*, *Cbx2*, *Cbx5*, *Dusp4*, and *E2f6*) satisfied both criteria (Figure 3D). *Arl4d* and *E2f6* have been identified as potential *let-7* target genes (Johnson et al., 2007, Li et al., 2015), suggesting that this screening successfully extracted candidate genes. Three of the five genes (*Cbx2*, *Cbx5*, and *E2f6*) are *Polycomb* group genes or *Polycomb*-associated genes (Core et al., 1997, Nielsen et al., 2001, Courel et al., 2008), suggesting that *Cbx5* and *E2f6*, as well as *Cbx2*, might be involved in *Hox* gene dysregulation via histone modifications and chromatin structural changes in *Lin28a*^-/-^ mice. Based on this screening, we examined if those five genes are true targets of *let-7* by using Luciferase assay. We generated reporter construct of luciferase-*let-7* target site mutated 3’UTR sequence of each gene and quantified *let-7* dependent reporter activity in comparison with Luciferase-wild type 3’UTR sequence construct (Figure 3E). We found that three of the five genes, *Cbx2*, *Cbx5,* and *E2f6*, were down-regulated in *let-7* dependent manner, whereas this down-regulation effect was not observed in *let-7* target site mutated construct. These data suggest that these genes are direct targets of *let-7* (Figure 3F).

To confirm that these genes are affected by *Lin28*/*let-7* axis *in vivo*, we performed mRNA and protein expression analyses on somite and neural tubes. qPCR analyses showed that *Arl4d* was significantly downregulated in *Lin28a*^-/-^ embryos. However, luciferase assay revealed that *Arl4d* expression did not affect *let-7* target site mutation, suggesting that *Arl4d* is not a direct target of *let-7* (Figure 3G). Protein expression analyses revealed that *Cbx2* was the only gene that was significantly downregulated in *Lin28a*^-/-^ embryos, and also its expression was affected in a *let-7*-dependent manner (Figure 3F and 3H). These findings suggest that *Cbx2* is, at least in part, a molecular target of the *Lin28a*/*let-7* pathway in skeletal patterning.

To support this concept further, we generated *Cbx2* mutant mice using CRISPR/Cas9 and interbred the *Cbx2* mutant with *Lin28a*^+/–^ mice (Figure S5A). The *Cbx2* homozygous mutants exhibited homeotic transformations (Figures S5B–S5E). Similar observations were reported for Cbx2-null mice (Core et al., 1997, Katoh-Fukui et al., 1998). We generated double heterozygous mutants of *Lin28a* and *Cbx2* (*Lin28a*^+/–^; *Cbx2*^+/–^) and analyzed their skeletal patterning. The double heterozygous mice showed ablation or truncation of the 13^th^ pair of ribs, although the *Lin28*^+/–^ and *Cbx2*^+/–^ single mutants did not show any obvious phenotypic irregularities (Figures 3I and 3J). These observations indicate that *let-7* directly regulates *Cbx2,* and that a genetic interaction exist between *Lin28a* and *Cbx2*. Taken together, these results suggest that the *Lin28a*/*let-7* reciprocal feedback regulates Cbx2 expression, and that this pathway is required for skeletal patterning during embryogenesis.

### *Lin28a*/*let-7* pathway modulates PRC1 occupancy at posterior *Hox* loci

*Hox* gene expression is epigenetically restricted to unique spatiotemporal patterns during embryogenesis by PcG genes (Soshnikova and Duboule, 2009). To determine if the *Lin28a*/*let-7*/*Cbx2* axis regulates skeletal patterning via *Hox* gene expression, we analyzed histone modifications and PcG occupancy at the *Hox* loci. We performed chromatin immunoprecipitation (ChIP) and q-PCR analyses on E9.5 somites and neural tubes (Figure 4A). For each assay, ChIP was performed on a pool of dissected somites and neural tubes from ten embryos (as n = 1). First of all, we analyzed the repressive histone modification (H3K27me3) at *Hoxa* cluster loci of Wt and We found the promoter regions of *Hoxa3*, *Hoxa9*, *Hoxa10*, *Hoxa11*, and *Hoxa13* exhibited a high concentration of histone H3K27me3 (Figure 4B). Intriguingly, the same genes were upregulated in *Lin28a*^−/−^ mice (Figure 2A), suggesting that these genes are tightly regulated epigenetically and that the loss of repressive histone modifications leads to the upregulation of these *Hox* genes.

**Figure 4.**
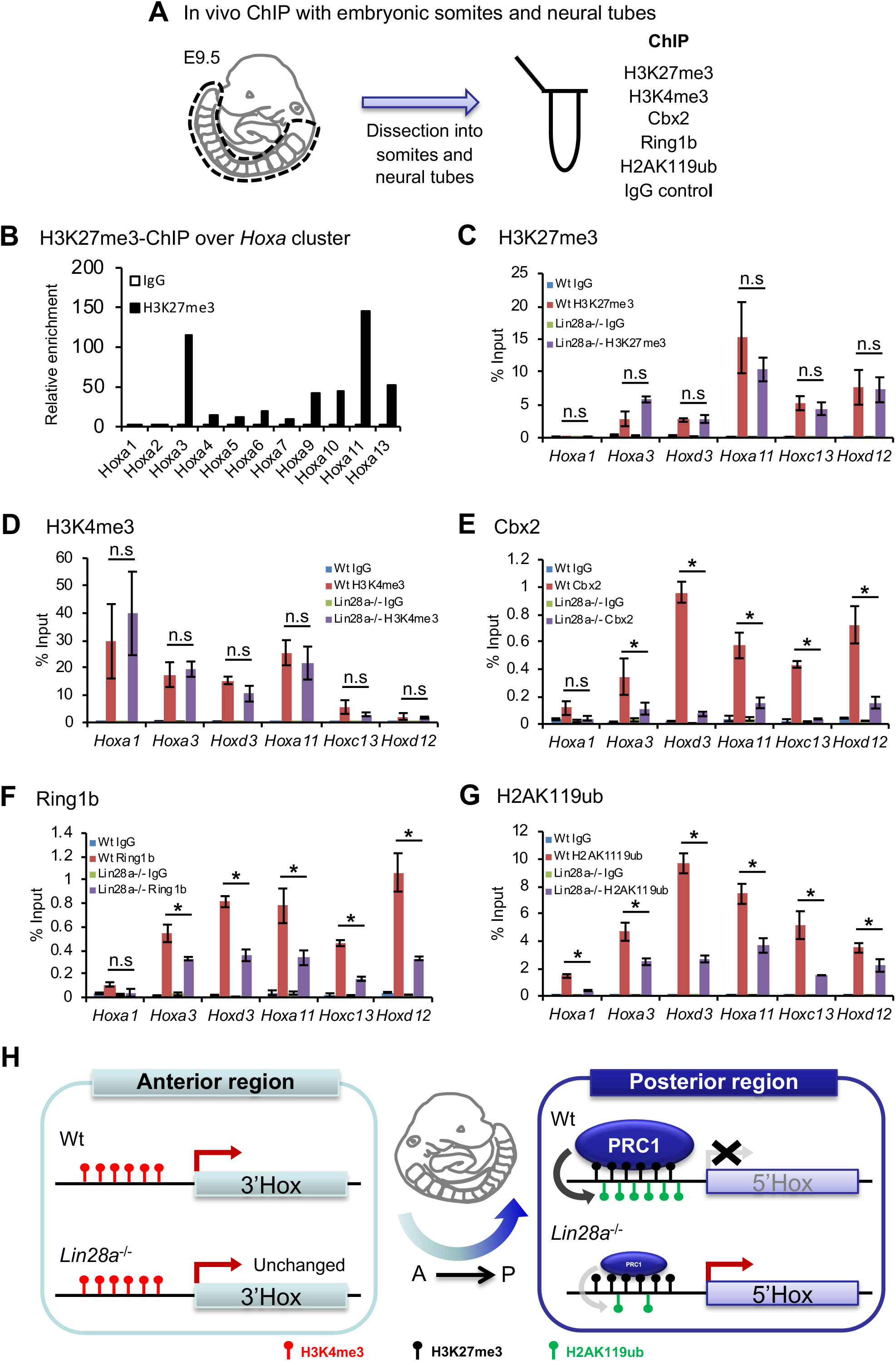
Histone modifications and polycomb occupancy at *Hox* loci in *Lin28a*^-/-^ **mice**. (**A**) Schematic diagram of the experimental procedure for ChIP analysis. (**B**) ChIP and q-PCR analyses of H2K27me3 in *Hox* A cluster genes in Wt embryos. (**C–G**) ChIP and q-PCR analyses of H3K27me3 (C), H3K4me3 (D), Cbx2 (E), Ring1b (F), and H2AK119ub (G). Percentages of immunoprecipitated DNA compared with the input are shown. (**H**) Summary of the chromatin state of *Hox* loci in Wt and *Lin28a*^−/−^ embryos. All data are expressed as the mean ± SEM (n = 3). * *p* < 0.05. n.s., not significant.

Based on the analysis of phenotype of the *Lin28a*^−/−^ homeotic transformation (Figure 1B-I) and *Hox* cluster gene expression pattern of *Lin28a*^−/−^ embryo (Figure 2A-C), we focused on *Hoxa3* and *Hoxd3* which are involved in C1-C2 malformation and partial fusion in knockout mice (Condie and Capecchi, 1994). In addition, we focused on *Hoxa11* which has been reported as the responsible gene for T13 to L1 homeotic transformation in mutated mice (Small and Potter, 1993), *Hoxd12* and *Hox*c*13* which were upregulated in *Lin28a*^−/−^ embryos, and *Hoxa1* as a representative of the anterior *Hox* gene (Figures 4C-G). Subsequently, we then performed ChIP and q-PCR analyses using anti-H3K27me3 and anti-H3K4me3 antibodies in Wt and *Lin28a*^−/−^ embryos (Figures 4C and 4D). We found that for histone H3 modifications, both K27me4 and K4me3 were not altered in *Lin28a*^−/−^ embryos compared with Wt (Figures 4C and 4D). We also performed ChIP using antibodies against PRC1 components to test their occupancy at *Hox* loci (Figures 4E and 4F). Consistent with the expression level of Cbx2 (Figure 3G), we found a two-fold reduction of its binding at posterior *Hox* regions in *Lin28a*^−/−^ mice (Figure 4E). Intriguingly, the occupancy of Ring1b, another component of PRC1 (Suzuki et al., 2002), and H2AK119 ubiquitination (H2AK119ub) which is catalyzed by Ring1b (Suzuki et al., 2002) also reduced in *Lin28a*^−/−^ mice (Figure 4F-G). Because each posterior *Hox* gene (*Hoxa11*, *Hoxc13*, and *Hoxd12*) is located on distinct chromosomes, these results indicate a critical role for the *Lin28a*/*let-7* axis in PcG-mediated *Hox* gene repression. Taken together, these findings suggest that Cbx2 repression by *let-7* leads to the reduction of PRC1 occupancy at the *Hox* loci and the transcriptional initiation of posterior *Hox* genes (Figure 4H).

### *Let-7* knockdown rescues *Hox* gene dysregulation in *Lin28a*^−/−^ cells

To further elucidate the importance of the direct regulation of *let-7* by Lin28a during *Hox* gene regulation, we tested whether *Hox* gene dysregulation could be rescued by knockdown of *let-7*-family microRNAs. To accomplish this, *Lin28a*^−/−^ embryonic stem (ES)-like cells were established from mutant blastocysts. Each *Lin28a*^−/−^ clone resembled Wt cells (Figure 5A), and we confirmed that the Lin28a protein was not detected in *Lin28a*^−/−^ ES cells (Figure 5B). These colonies showed high alkaline phosphatase activity (Figure 5A) and also expressed pluripotent factors (Figure 5C). As observed in *Lin28a*^−/−^ embryos (Figure 3B), global accumulation of *let-7*-family microRNAs was observed in the mutant cells (Figure 5D).

**Figure 5.**
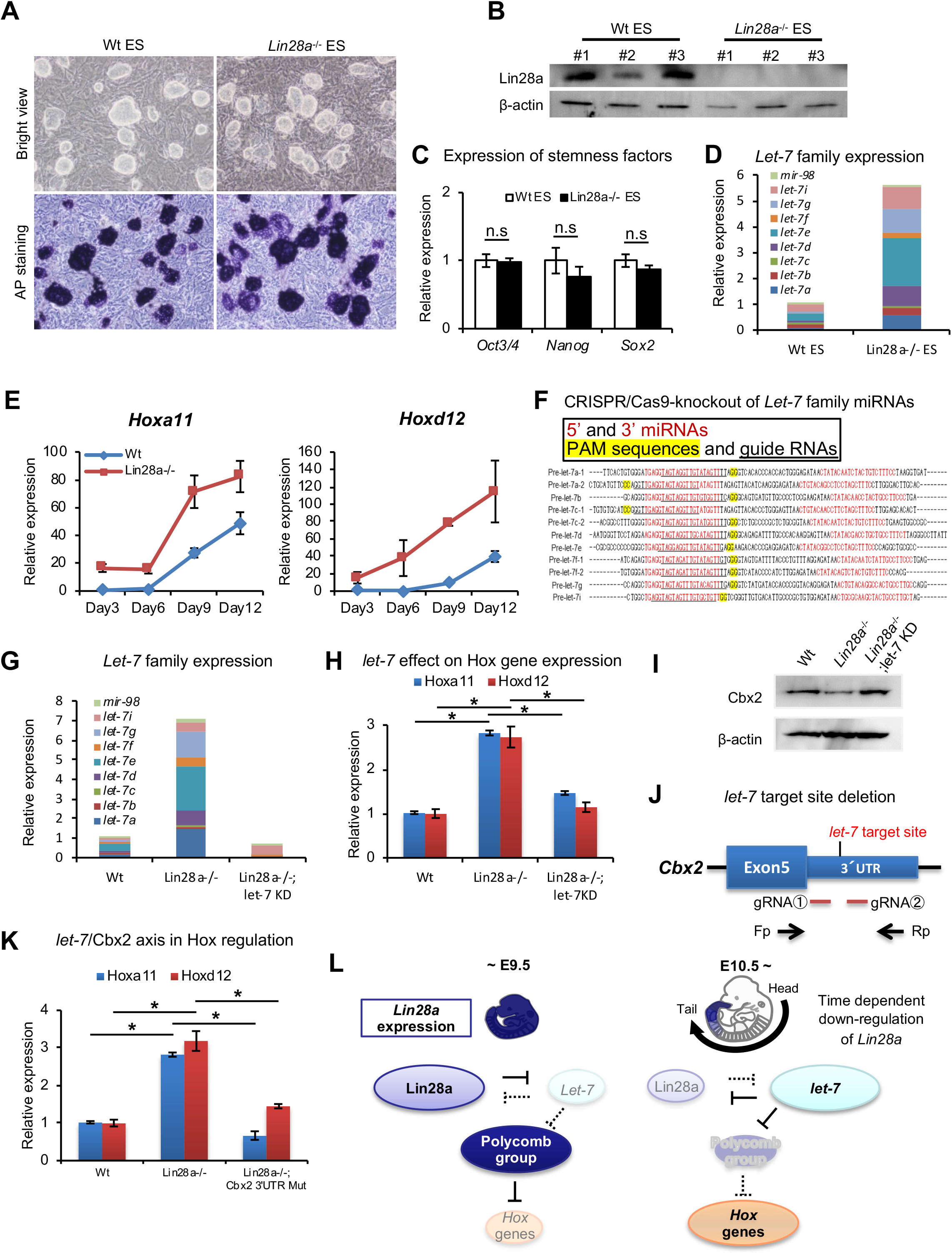
Knockdown of *let-7* can reverse *Hox* gene dysregulation. (**A**) Morphology (top panels) and alkaline phosphatase activity (bottom panels) of Wt and *Lin28a*^−/−^ ES-like cells. (**B**) Western blot analysis of Lin28a in ES-like cells. β-actin is shown as a loading control. (**C**) q-PCR analysis of stemness factors. (**D**) q-PCR analysis of *let-7*-family members. The level of expression relative to total let-7 amount in Wt is shown. (**E**) q-PCR analyses of *Hoxa11* and *Hoxd12* over a time course of 3, 6, 9, 12 days following embryoid body formation. (**F**) Precursor sequences of *let-7*-family members and guide RNAs for *let-7* targeting Let-7 mature microRNAs are shown in red. The protospacer adjacent motif (PAM) sequence for hCas9 is highlighted in yellow, and targeting sequences are underlined. (**G**) *Let-7* expression in Wt, *Lin28a*^−/−^ and *Lin28a*^−/−^; *let-7* KD cells. The level of expression relative to total let-7 amount in Wt is shown. (**H**) *Let-7* knockdown rescues *Hox* gene dysregulation in *Lin28a*^−/−^ cells. (**I**) Cbx2 expression level of Wt, *Lin28a*^−/−^ and *Lin28a*^−/−^; *let-7* KD derived EBs. β-actin is shown as a loading control. (**J**) Schematic diagram of *let-7* target site deletion from *Cbx2* 3’UTR and genotyping via PCR of mutant clones. (**K**) q-PCR analyses of *Hoxa11* and *Hoxd12* following embryoid body formation. (**L**) Schematic diagram of *Lin28a*/*let-7* mediated *Hox* gene regulation. All data are expressed as the mean ± SEM (n = 3). n.s., not significant.

In the following experiments, we differentiated ES cells to embryoid bodies. ES cells and embryoid bodies require different PRC1 components to maintain their state. ES cells are maintained in an undifferentiated state, using Cbx7 containing PRC1. On the other hand when ES cells exit the pluripotent state and differentiate into embryoid bodies, Cbx2 is expressed and becomes a component of PRC1 (Morey et al., 2012). Thus, we utilized embryoid bodies as an appropriate model to analyze *Hox* genes via *Lin28*/*Let-7*/*Cbx2* axis. Embryoid bodies were produced from each clone and expression changes of *Hox* genes were analyzed. *Hox* genes were upregulated upon differentiation in these embryoid bodies, suggesting that a recapitulation of the *Hox* gene upregulation observed in *Lin28a*^−/−^ mice occurred in *Lin28a*^−/−^ ES-like cells (Figure 5E).

Next, we knocked down the *let-7* family in *Lin28a*^−/−^ ES-like cells using the CRISPR/Cas9 system to test if *Hox* gene upregulation could be rescued by the reduction of *let-7* microRNAs. The major *let-7* family is composed of 11 genes (*a-1*, *a-2*, *b*, *c-1*, *c-2*, *d*, *e*, *f-1*, *f-2*, *g*, and *i*), and we performed the knockdown of this series of *let-7* genes using guide RNAs targeting *let-7s* (Figure 5F). The clone that yielded a highly efficient deletion of *let-7* microRNAs in *Lin28a*^−/−^ cells (*Lin28a*^−/−^; *let-7*KD) was selected for further analyses. We confirmed the accumulation of *let-7* in *Lin28a*^−/−^ cells and the drastic reduction in *Lin28a*^−/−^; *let-7*KD clones (Figure 5G). qPCR analysis revealed that dysregulation of Hoxa11 and Hoxd12 was rescued in *Lin28a*^−/−^; *let-7*KD clones (Figure 5H). Moreover, we also confirmed that a decreasing Cbx2 protein expression level in embryoid bodies-derived *Lin28a*^−/−^ ES-like cells was rescued by knock down of *let-7* (Figure 5I).

To directly prove that the *Lin28a*^−/−^ phenotype results from the *let-7*-mediated down-regulation of *Cbx2*, we established *Lin28a*^-/-^ ES cells with *let-7* target site deletion from *Cbx2* 3’UTR (*Lin28a*^−/−^; *Cbx2* 3’UTR mutant) using CRISPR/Cas9 system, and we examined whether *let-7* target site deletion from *Cbx2* 3’UTR could rescue *Hox* gene dysregulation. Two guide RNAs targeting *let-7* binding site in *Cbx2* 3’UTR were constructed and were transfected with Cas9 expression vector into *Lin28a*^−/−^ ES-like cells for the establishment of *Cbx2* 3’UTR mutant cell lines (Figure 5J). Furthermore, we generated embryoid bodies from Wt, *Lin28a*^−/−^, and *Lin28a*^−/−^; *Cbx2* 3’UTR mutant clones in the same manner as that of *let-7* knock down experiment (Figure 5G-I), and the expression level of *Hox* genes were analyzed. Consistent with the result of *let-7* knock down (Figure 5G-I), we found that *Hoxa11* and *Hoxd12* were up-regulated in *Lin28a*^−/−^ cells, and that this abnormal expression was absent in *Lin28a*^−/−^; *Cbx2* 3’UTR mutant cells (Figure 5K). These results suggest that *let-7*-mediated Cbx2 repression might be, at least in part, responsible for *Hox* gene dysregulation in Lin28a^−/−^ mice. Taken together, our results suggest that the upregulation of *let-7* leads to decreased PRC1 occupancy, which causes the disruption of the “*Hox* code,” thus indicating the potential role of *Lin28a*/*let-7* pathway in skeletal patterning via *Polycomb*-mediated *Hox* gene regulation (Figure 5L).

## Discussion

The body plan along the anteroposterior axis is tightly regulated by *Hox* genes. During development, each *Hox* gene must be activated at a precise position with precise timing. Spatiotemporal regulation via chromatin conformational changes is essential for *Hox* gene expression and for subsequent anteroposterior patterning (Soshnikova, 2014); however, the molecular mechanisms behind these processes are not fully understood. In this study, we demonstrated the fundamental role of *Lin28a*/*let-7* pathway in skeletal patterning and vertebral specification. *Lin28a*-mediated repression of *let-7* biogenesis is required for *Cbx2* expression and *Hox* gene repression by PcG genes. It is known that the deletion mutants of the *Hox* early enhancer exhibit anterior transformations of vertebrae because of the heterochrony of *Hox* gene expression (Juan and Ruddle, 2003). In our *Lin28a*^−/−^ mice, posterior transformations were observed in the thoracic region (Figures 1D–1I), suggesting that developmental timing of *Hox* gene initiation occurs earlier than in Wt mice. Consistent with this speculation, precocious expression of *Hoxc13* causes premature arrest of axial extension, similar to that of *Lin28a*^−/−^ mice (Young et al., 2009, Mallo et al., 2010). This indicates that the tail truncation observed in *Lin28a*^−/−^ mice might be caused by spatiotemporal dysregulation of *Hoxc13* (Figures 2B and 2C). These observations of temporally diminished expression patterns of *Lin28a* during embryogenesis (Figure 1A) (Yokoyama et al., 2008, 2009) strongly support our hypothesis that the down-regulation of Lin28a triggers *let-7* expression, PRC1 displacement at *Hox* loci, and subsequent *Hox* gene activation in a developmental time-dependent manner (Figure 5L). The other *Hox* genes, including *Hoxa3*, *Hoxd3*, *Hoxb8*, *Hoxc8*, *Hoxa11*, and *Hoxa13*, showed no obvious difference in their expression patterns, although the expression of the genes were up-regulated (Figure 2A, S3). Homeotic transformations are mainly caused by the altered expression pattern of *Hox* gene, while it is also possible that homeotic transformation are caused altered the expression level of the *Hox* gene in each somite. For instance, *Dll1* enhancer driven *Hoxb6* transgenic mice show ectopic rib-like structure in cervical, lumber, sacral and caudal regions. However, malformation of axial skeleton are shown even in thorax which is the originally expressing region of *Hoxb6* (Vinagre et al., 2010). These results suggest the possibility that the *Hox*-code of the specific region might have been edited due to the specific *Hox* gene with elevated expression, which might cause the morphological change of vertebrae in our *Lin28a*^−/−^ mice.

PcG genes are regulators of the “*Hox* code” at the level of chromatin structure, which occurs via epigenetic histone modifications (Mallo and Alonso, 2013, Soshnikova, 2014). In ES cells, *Hox* genes are silenced in a bivalent state containing both H3K27me3, a repressive, and H3K4me3, an active histone marker. During development, the epigenetic status of *Hox* loci is dynamically balanced by PcG genes and *Trithorax* group (TrxG) genes, which are required for the trimethylation of H3K4. PcG genes should be repressed prior to the initiation of *Hox* gene expression to open the chromatin along the anteroposterior axis. However, the precise molecular mechanisms underlying the inhibition of the expression of PcG genes during embryogenesis are not fully understood. Here, we provide evidence that *Lin28a*/*let-7* pathway is, at least in part, one of the mechanisms that are involved in the regulation of PcG genes (Figure 3). Cbx2 is required for the binding of PRC1 to target loci and recognition of H3K27me3, and these processes are catalyzed by Ezh2, the main component of PRC2. *Ezh2* is directly targeted by *let-7* microRNAs in primary fibroblasts and cancer cells (Kong et al., 2012). In contrast with those findings, there were no apparent differences in the level of H3K27me3 at *Hox* loci in *Lin28a*^−/−^ mice (Figure 4C). *Ezh2*^−/−^ embryos died at the peri-and post-implantation stages (O’Carroll et al., 2001), whereas mutant mice of the PRC1 genes exhibited skeletal transformations (van der Lugt et al., 1994, Akasaka et al., 1996, Core et al., 1997, Suzuki et al., 2002, Li et al., 2011, Katoh-Fukui et al., 1998) that were similar to those of *Lin28a*^−/−^ mice (Figures 1D–1I). These observations suggest that *Lin28a*/*let-7* pathway is involved in the later phases of epigenetic silencing of *Hox* genes during skeletal patterning. We confirmed the genetic interaction between *Lin28a* and *Cbx2* (Figure 3I), and the reduction of PRC1 occupancy at *Hox* loci in *Lin28a*^−/−^ mice (Figures 4E and 4F). These findings indicate that *let-7*-mediated Cbx2 repression leads to the reduction of PRC1 occupancy at *Hox* loci, resulting in the transcriptional initiation of posterior *Hox* genes (Figure 4H).

In addition to epigenetic regulation by PcG genes, posttranscriptional regulation by microRNAs is also required for anteroposterior patterning. During mouse embryogenesis, mesoderm-specific ablation of Dicer, which is an RNase III enzyme that is required for microRNA biogenesis, results in a posterior shift in hindlimb position (Zhang et al., 2011), suggesting the involvement of microRNAs in normal skeletal patterning and vertebrae specification. Two microRNA families, *mir-10s* and *mir-196s*, are located in *Hox* clusters, and they are thought to regulate *Hox* gene expression and specify the regional identities along the anteroposterior axis (Heimberg and McGlinn, 2012). It has also been reported that the *mir-17-92* cluster, which contains *mir-17*, *mir-18*, *mir-19*, *mir-20*, and *mir-92*, is required for normal skeletal patterning (Han et al., 2015). Although *Lin28a* is a regulator of microRNA biogenesis, the expression of these microRNAs was not altered in the *Lin28a*^−/−^ mice compared with Wt animals (Figures 3A and 3C). These results suggest that *Lin28a/let-7* pathway acts independently of these microRNAs in *Hox* gene regulation. *Mir-10s* and *mir-196s* are involved in the spatial regulation of *Hox* genes to shut down target *Hox* genes in specific regions (Heimberg and McGlinn, 2012), whereas *let-7* might be required for temporal activation of Hox genes via Lin28a downregulation during development. These results suggest that *let-7* can be distinguished from other microRNAs in skeletal patterning, and that *Lin28a*/*let-7* pathway links posttranscriptional regulation to PcG-mediated epigenetic regulation in *Hox* gene regulation.

MicroRNAs are thought to regulate hundreds of target genes and to modulate multiple biological processes, and hence, the accumulation of *let-7* observed in *Lin28a*^−/−^ mice might lead to extensive disorders of gene regulatory networks. It is well known that the *let-7* family regulates *c-Myc*, *K-ras*, *Hmga2*, and other genes that are involved in cell proliferation and oncogenesis (Mayr et al., 2007, Lee and Dutta, 2007, Johnson et al., 2005, Sampson et al., 2007). Knockout mice for these genes exhibit dwarfism caused by a reduction of cell proliferation that is similar to that observed in *Lin28a*^−/−^ mice (Zhou et al., 1995, Koera et al., 1997, Johnson et al., 1997, Trumpp et al., 2001). These observations suggest that the growth defects and postnatal mortality of *Lin28a*^−/−^ mice (Figure S2) may be attributed to the dysregulation of such genes; however, their requirement for skeletal patterning has not been characterized. Despite the contribution of these genes to the *Lin28a*^−/−^ phenotype, it is noteworthy that there was a genetic interaction between *Lin28a* and *Cbx2* during skeletal patterning (Figure 3I). These results suggest that the *Lin28a*/*let-7*/*Cbx2* pathway is, at least in part, responsible for normal skeletal patterning. In addition to *Lin28a, Lin28b* regulates *let-7* biogenesis, and it is known that single nucleotide polymorphisms (SNPs) of the human *LIN28B* locus correlated with height and the timing of menarche (Lettre et al., 2008, Perry et al., 2009, He et al., 2009, Ong et al., 2009, Widen et al., 2010, Sulem et al., 2009). These studies suggest that the regulation of developmental timing by *Lin28b* is also conserved in mammals; however, its requirement in skeletal patterning is still unclear.

Based on the analysis of phenotype of the *Lin28a*^−/−^ homeotic transformation (Figure 1B-I) and *Hox* cluster gene expression pattern of *Lin28a*^−/−^ embryo (Figure 2A-C), we focused on *Hox3a*/*d3* which are involved in C1-C2 malformation and partial fusion in knockout mice (Condie and Capecchi, 1994), *Hoxa11* which has been reported as the responsible gene forT13 to L1 homeotic transformation in mutated mice (Small and Potter, 1993), *Hox12d*/c*13* which is upregulated in *Lin28a*^−/−^ embryos, and *Hoxa1* as a representative of anterior *Hox* gene (Figures 4C-G).

We first examined histone H3K27me3/H3K4me3 modification and occupancy of Cbx2 and Ring1b at each *Hox* gene promoter region. We found that for histone modifications, both H3K27me3 and H3K4me3 were not altered in *Lin28a*^−/−^ embryos compared with Wt (Figure 4C-D). In contrast, we found a two-fold reduction of its binding at *Hox* gene promoter regions in *Lin28a*^−/−^ mice (Figure 4E), which was consistent with the expression level of Cbx2 (Figure 3H). Interestingly, the occupancy of Ring1b was also reduced in *Lin28a*^−/−^ mice (Figure 4F). Moreover, we analyzed the abundance of ubiquitination of H2A (H2AK119ub), which is catalyzed by Ring1b. We found that not only Ring1b occupancy but also H2AK119ub was decreased at *Hox* gene promoter. These data indicate that *Lin28a*^−/−^ phenotype correlated with PRC1 dysregulation especially Cbx2 dysregulation and strongly suggest the existence of a molecular pathway of *Lin28*/*let-7*/*Cbx2*. To further support our hypothesis, an alternative analysis to demonstrate the direct regulation of *Cbx2* via *Lin28*/*let-7* axis were carried out. We performed rescue experiment by the deletion of *let-7* target site from *Cbx2* 3’UTR *in Lin28a*^−/−^ ES cells (*Lin28a*^−/−^; *Cbx2* 3’UTR mutant). *Lin28a*^−/−^; *Cbx2* 3’UTR mutant ES cells were established using CRISPR/Cas9 system (Figure 5J). Cbx2 becomes a component of PRC1 only when ES cells exit the pluripotent state (Morey et al., 2012). Hence, embryoid bodies were generated from Wt, *Lin28a*^−/−^, and *Lin28a*^−/−^; *Cbx2* 3’UTR mutant ES cells, and the expression level of *Hox* genes were analyzed. Consistent with the result of *let-7* knock down in *Lin28a*^−/−^ ES cells-derived embryoid bodies (Figure 5H), *Lin28a*^−/−^; *Cbx2* 3’UTR mutant ES cells-derived embryoid bodies showed reduced expression levels of *Hoxa11* and *Hoxd12* which were comparable with those of Wt ES cells-derived embryoid bodies (Figure 5K). These data directly showed the regulation of *Cbx2* via *Lin28*/*let-7* axis and clearly confirmed the existence of *Lin28*/*let-7*/*Cbx2* molecular pathway.

The expression level of Cbx2 was also downregulated in heterozygous *Lin28a*^+/-^ (Figure S6). This may indicate that Lin28a expression in heterozygous *Lin28a*^+/-^ is strongly reduced to less than half (Figure S1C), suggesting that other target molecules regulated by Lin28a might be involved in this homeotic transformation phenotype. In addition to the regulation of *let-7*, it is also known that Lin28a and its homolog, Lin28b, bind to and modulate the translation efficiency of specific mRNAs, such as *Igf2*, *Oct4*, *Ccnb1*, *Cdk6*, *Hist1h2a*, and *Bmp4* (Xu et al., 2009, Ma et al., 2013, Qiu et al., 2010, Xu and Huang, 2009). Moreover, recent HITS-CLIP and PAR-CLIP technology identified a variety of mRNAs as *Lin28* family targets (Wilbert et al., 2012, Madison et al., 2013, Hafner et al., 2013, Cho et al., 2012). Among them, two studies showed that *Lin28* family might have a potential to bind specific *Hox* genes in HEK293T, DLD1, and Lovo cell lines (Lin28a to *Hoxa9*, *a11*, *b4*, *b6*, *b9*, *c4*, *d11*; Lin28b to *Hoxa9*, *b3*, *b4*, *b7*, *b8*, *b9*, *d13*) (Hafner et al., 2013; Madison et al., 2013), whereas CLIP-Seq analysis with ES cells did not show such possibility (Cho et al., 2012). Moreover, *Cbx5* is a *Lin28a* target gene as well as one of the potential *let-7* targets. *Cbx5* encodes heterochromatin binding protein, and the depletion of this gene causes skeletal defects in mice, although protein level of Cbx5 was not altered in *Lin28a*^−/−^ mice. These previous reports and our results imply that both *let-7*-dependent and -independent function of *Lin28a* might affect skeletal patterning during development. However, further studies are required to deepen the understanding of the developmental functions of *Lin28* family and its involvement in skeletal patterning.

Recently, two independent groups reported the function of Lin28 family as regulators of trunk elongation (Aires et al., 2019, Robinton et al., 2019). Tail bud specific overexpression of *Lin28* family increased caudal vertebrae number (Aires et al., 2019, Robinton et al., 2019). However, the loss of *Lin28* in tail bud resulted in the reduction of caudal vertebrae number (Robinton et al., 2019). These results are consistent with our *Lin28a* ^-/-^ mice phenotypes with short stature and shortened tails (Fig1B, FigS2A, B). Furthermore, Aires et al. showed that Lin28 and Hox13 had opposite function in tail bud proliferation, suggesting that the balance of the expression of those two genes, which might be regulated by GDF signaling, is one of the determinants of the tail length (Aires et al., 2019). Our results revealed the epigenetic inhibition of HoxPG13 by *Lin28a*/*let-7*/*Cbx2* pathway, which might be one of the mechanisms that explain the antagonistic function of Lin28a and HoxPG13 in axial elongation as well as in skeletal patterning. Robinton et al. showed that Lin28a regulated cell fate choice between mesodermal cells and neural cells; however, no homeotic transformations were observed in their *Lin28a* overexpression mutants. Further analyses are required to determine if the homeotic mutations found in *Lin28a*^-/-^ mice is caused by the dysregulation of cell fate choice.

Taken together, our results suggest that the negative feedback between *Lin28a* and *let-7* regulates the PRC1 component, *Cbx2,* and the subsequent spatiotemporal expression of *Hox* genes during mammalian embryogenesis. The loss of Lin28a caused homeotic transformations via the premature loss of PRC1 at the promoter region of posterior *Hox* genes, thus establishing a new role of *Lin28a*/*let-7* pathway in the modulation of the “*Hox* code.” It is of interest to test whether this role of *Lin28a*/*let-7* in *Hox* regulation was acquired in the evolutional process, or it has always been involved in heterochrony in *C. elegans*.

## Materials and methods

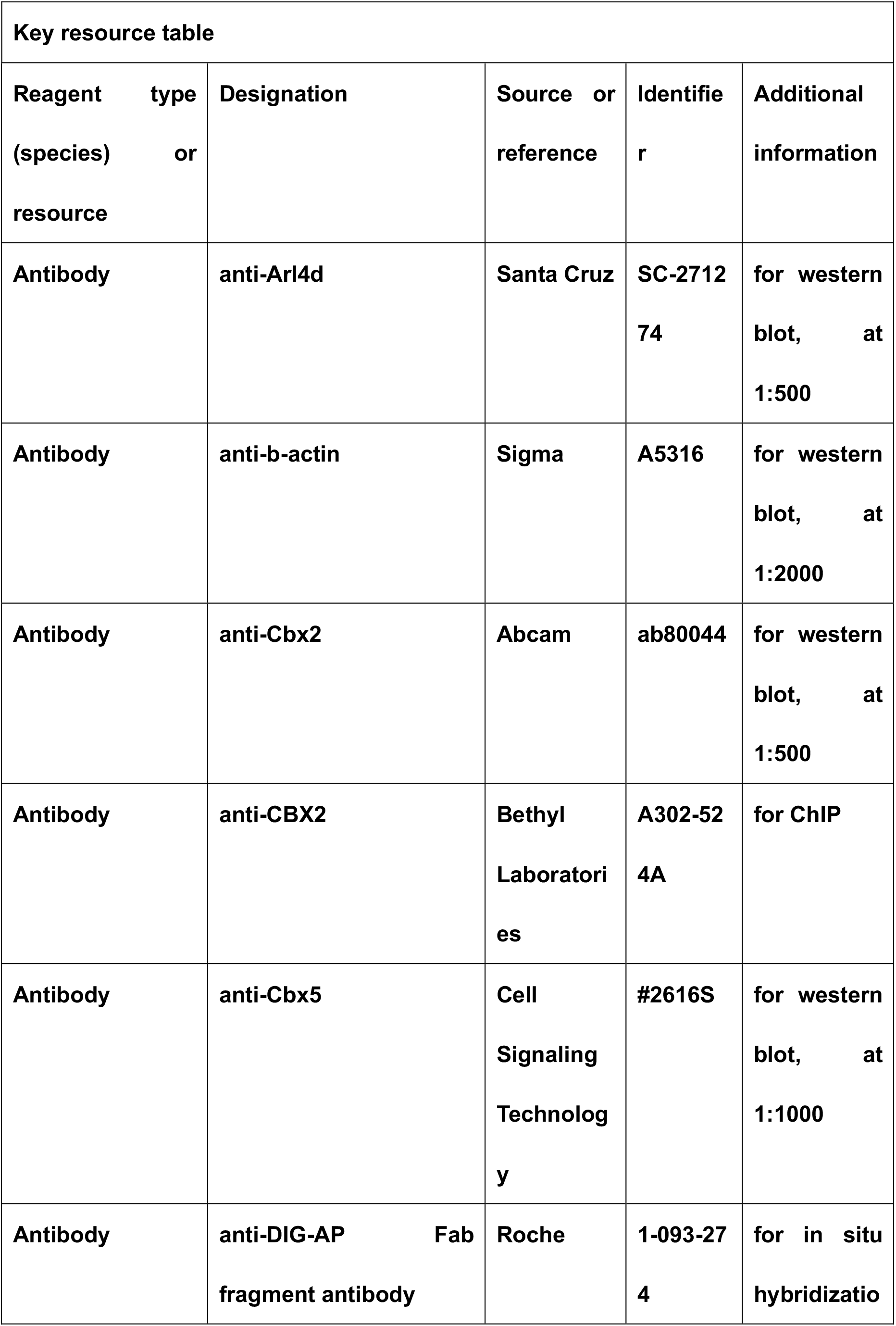

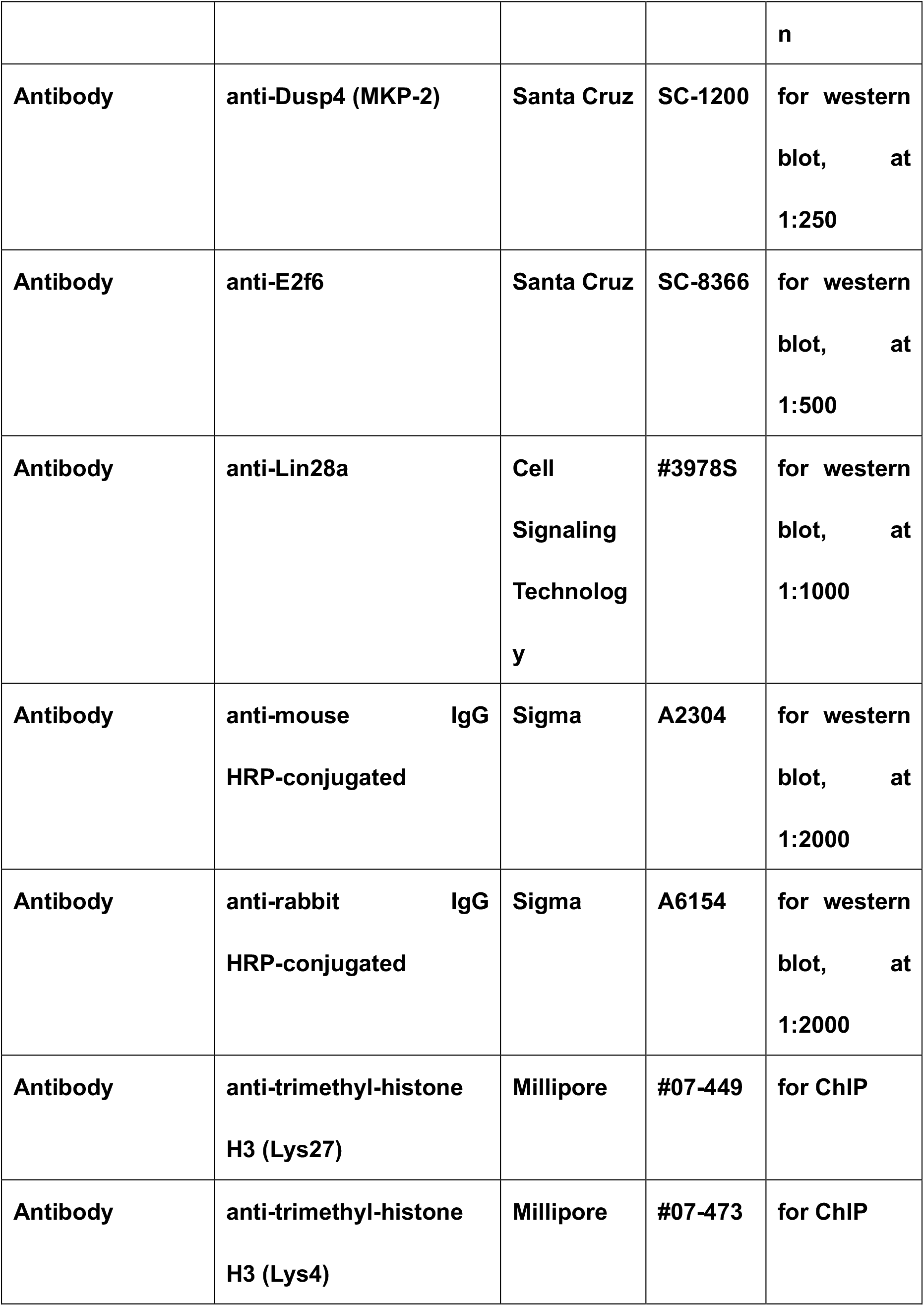

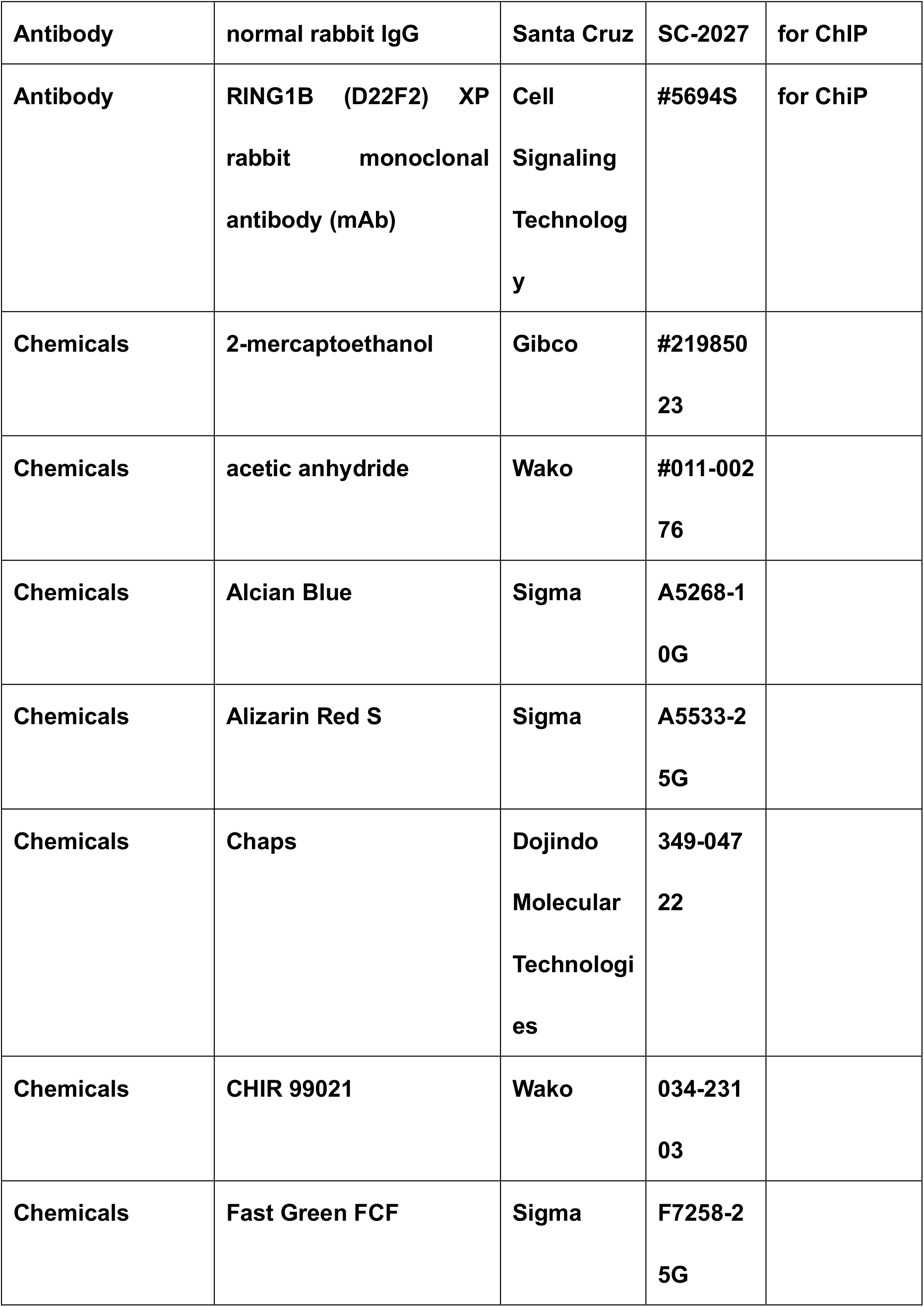

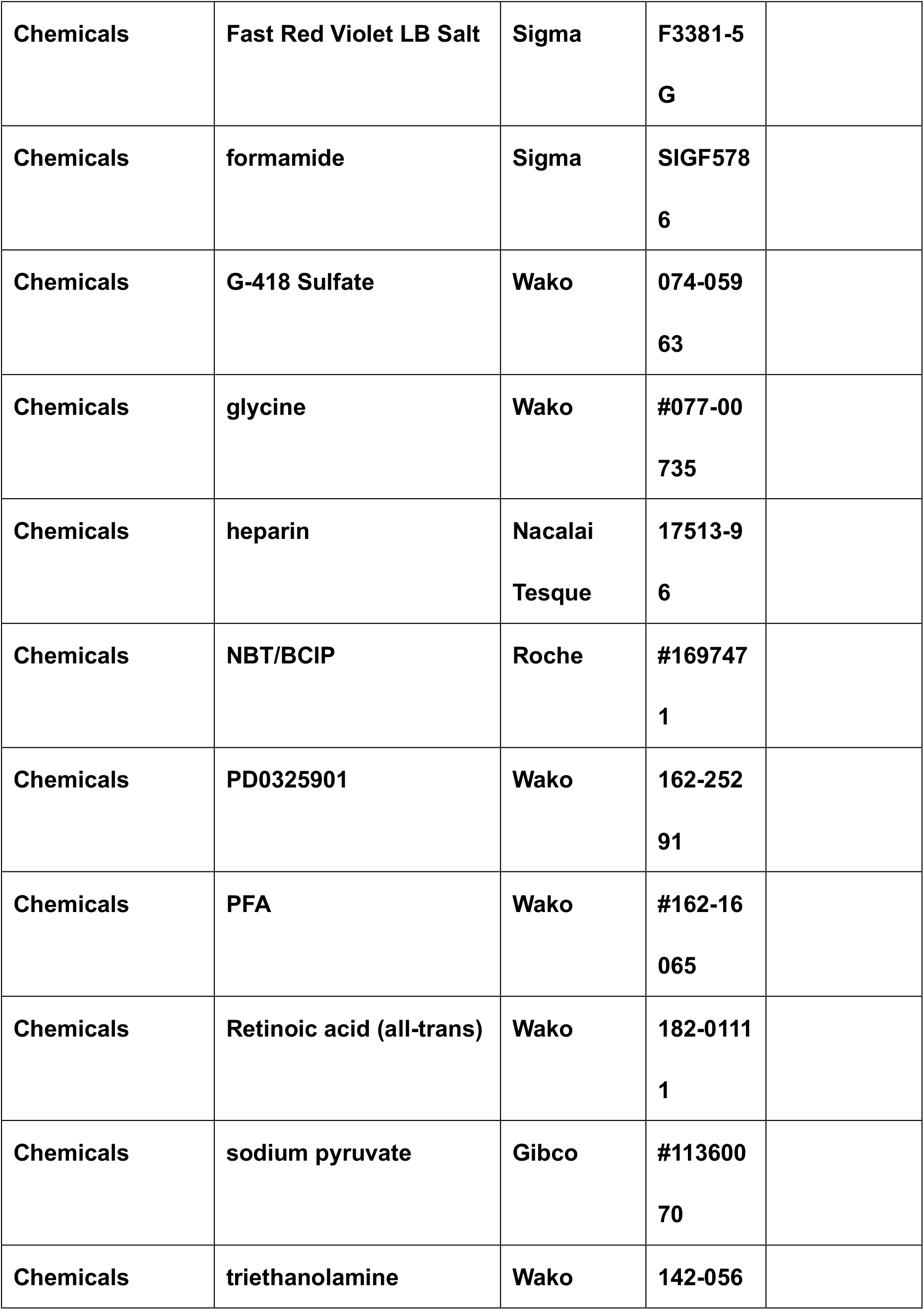

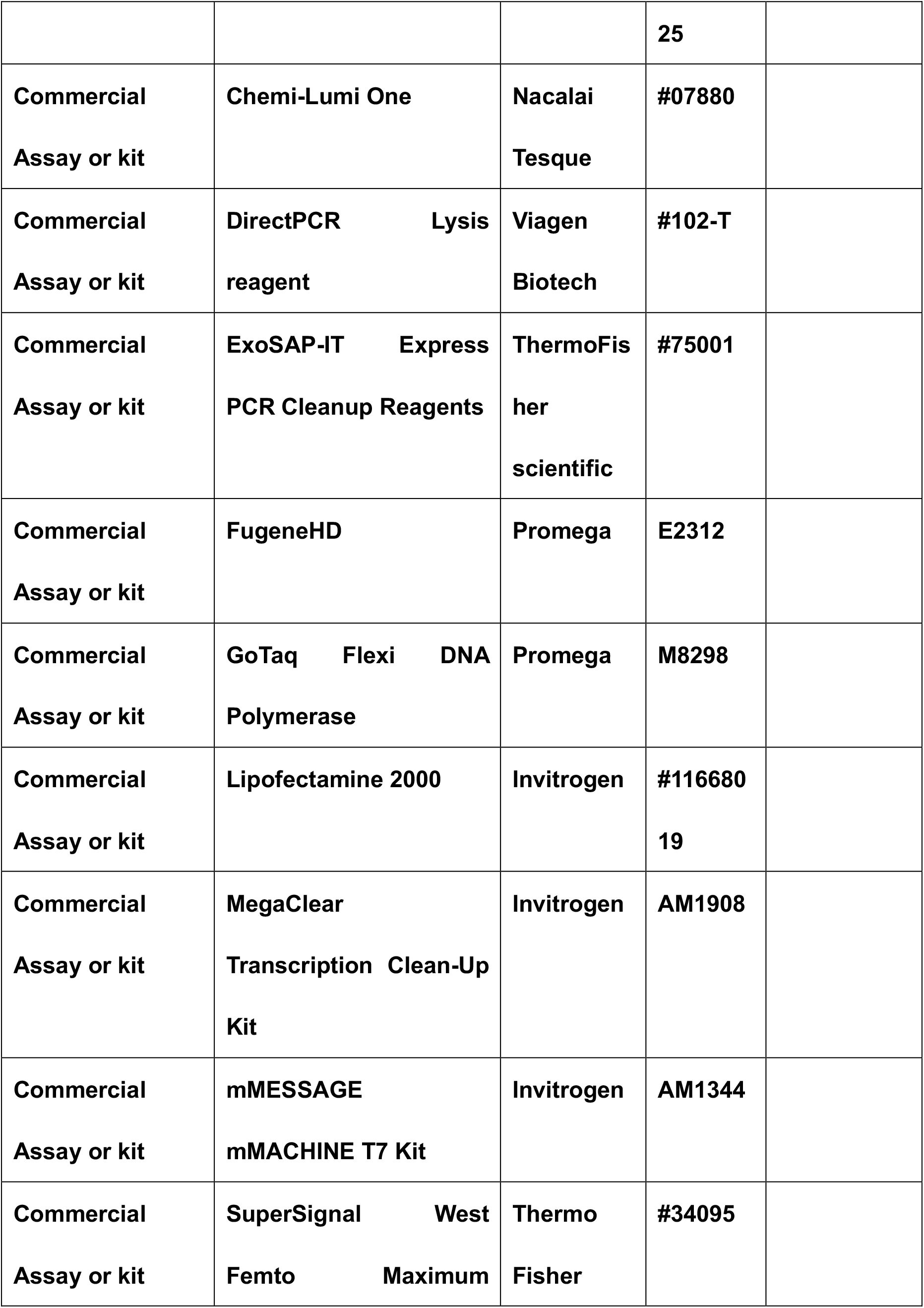

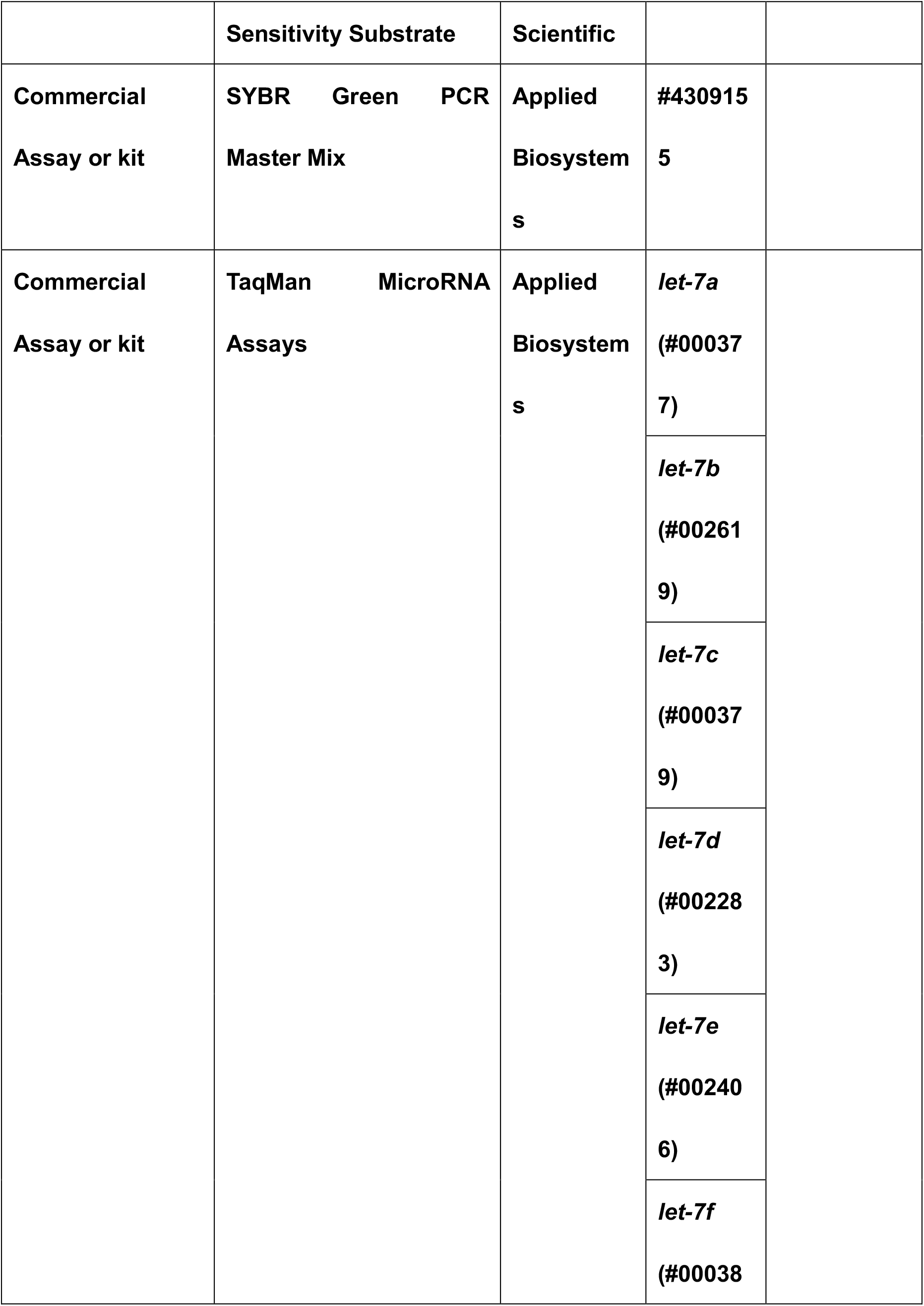

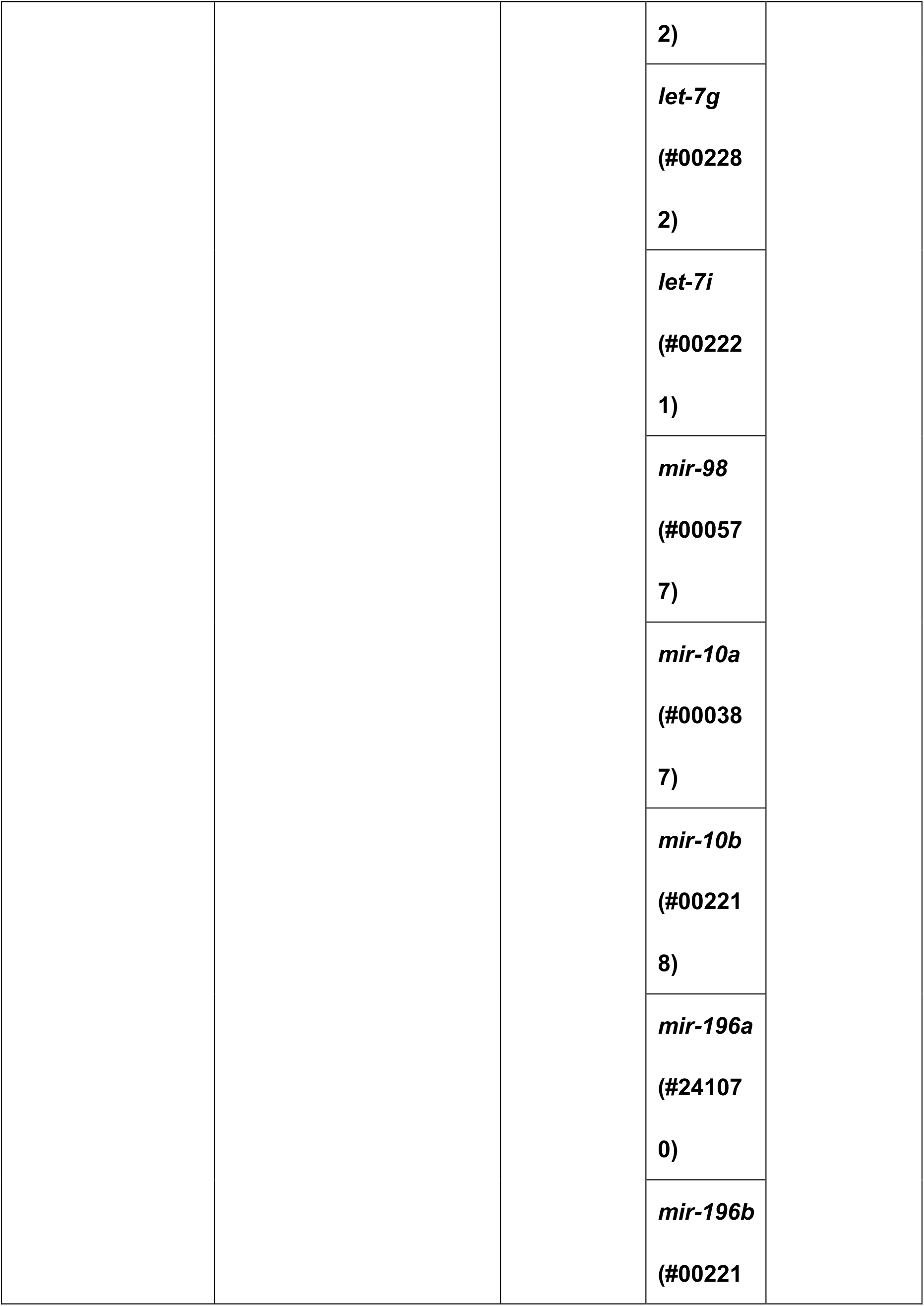

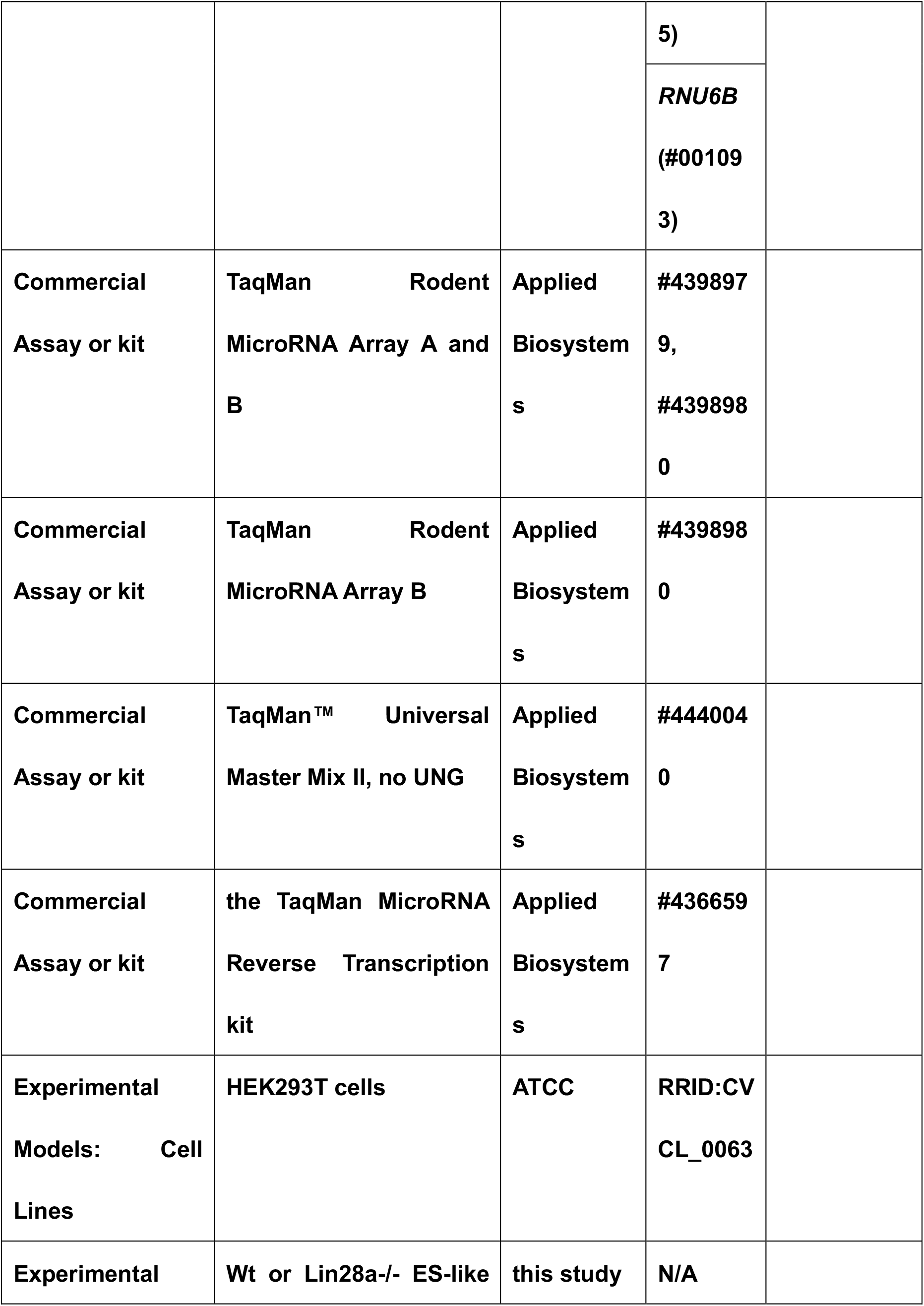

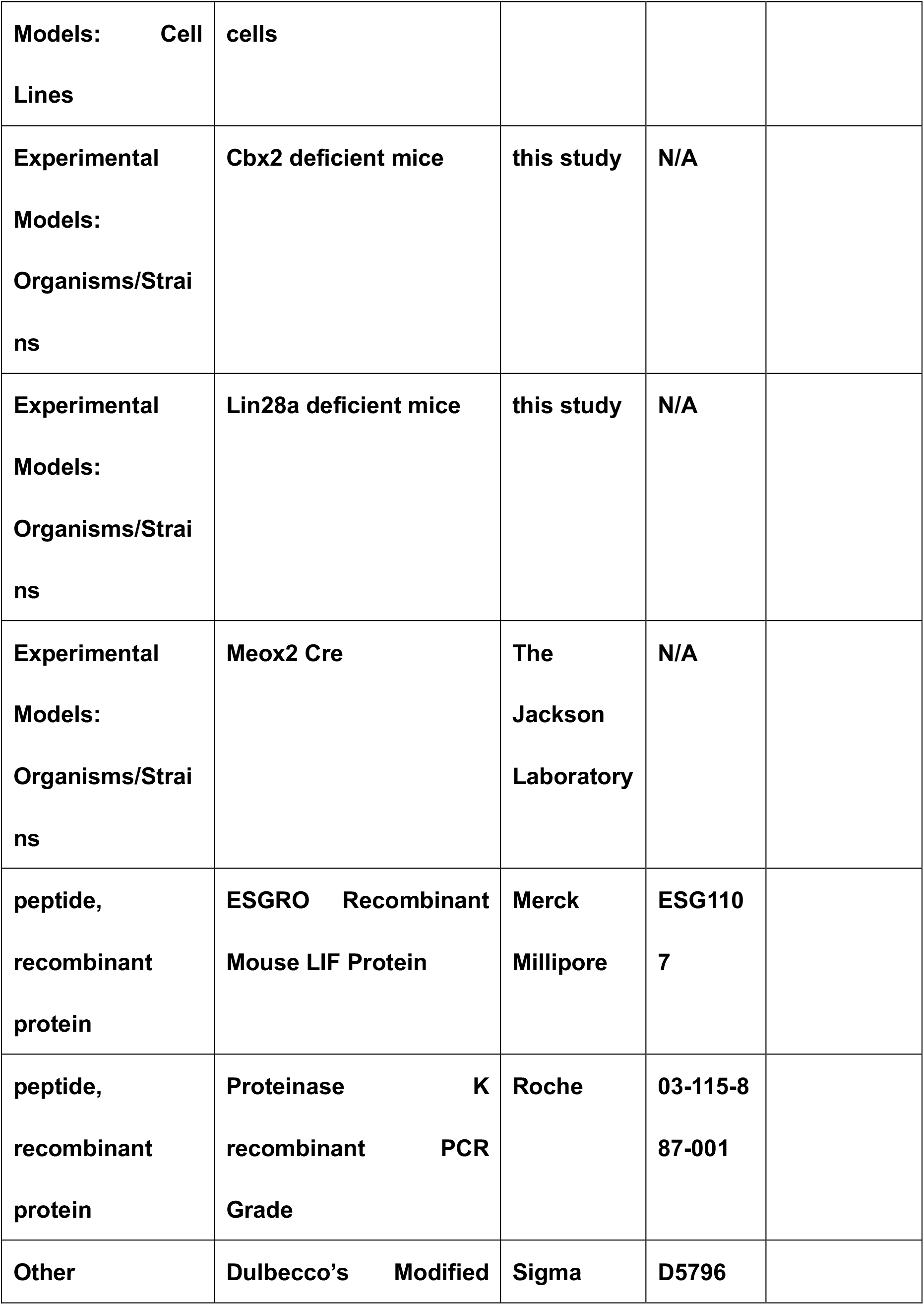

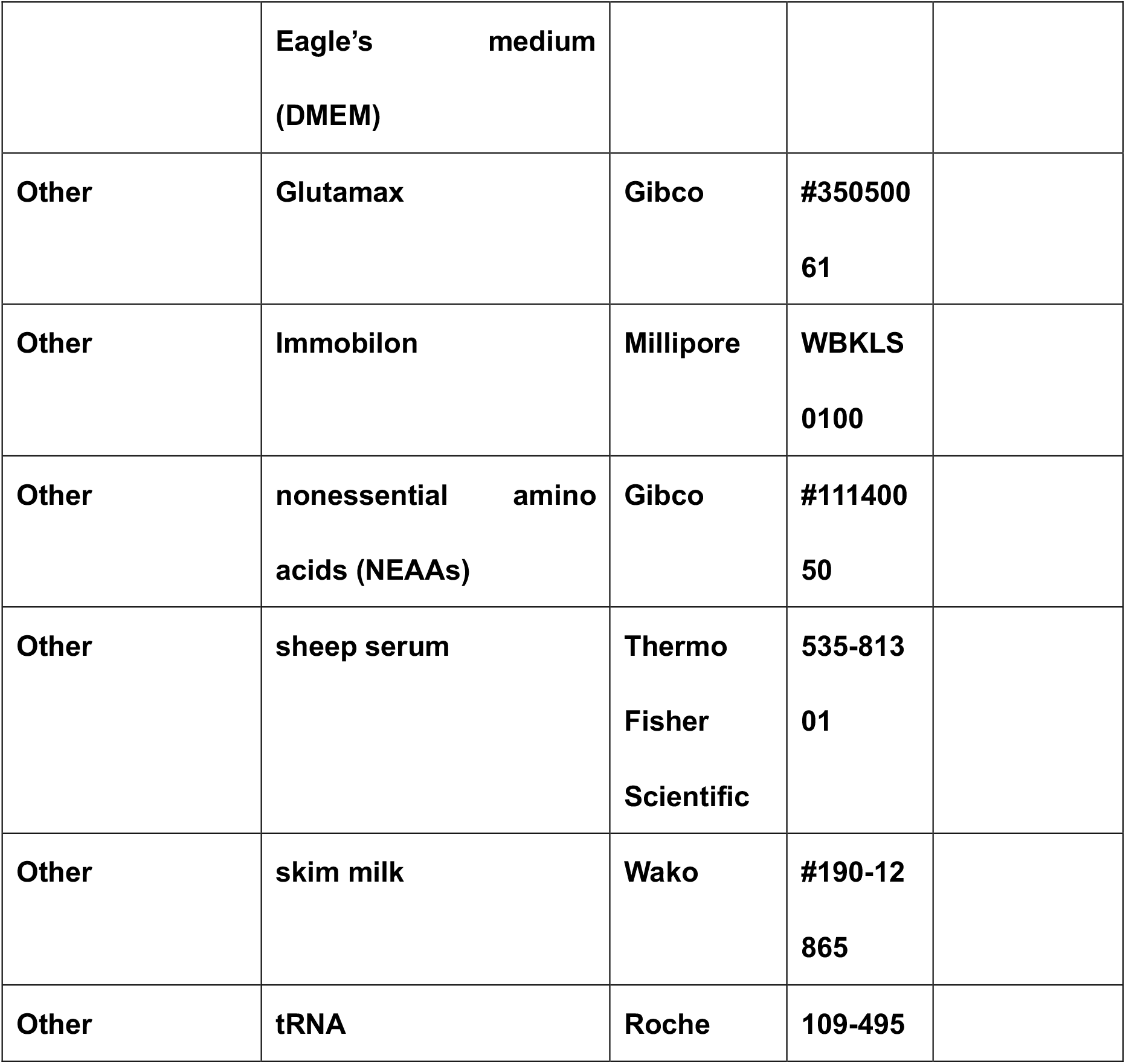

### Generation of mutant mice

All animal experiments were performed in accordance with protocols approved by the Institutional Animal Care and Use Committee of the National Research Institute for Child Health and Development (permit numbers: 2004-003, 2014-001). To accomplish the *Lin28a* knockout, the targeting vector was constructed to replace the endogenous *Lin28a* locus with the Venus gene and PGK-neo cassette by homologous recombination in ES cells. The 5′ and 3′ sequences flanking the endogenous *Lin28a* locus were amplified by PCR from a C57BL/6N genomic *bacterial artificial chromosome* (BAC) clone (BACPAC Resource Center). The primer sequences used for homology arm cloning were as follows: 5′ homology arm forward primer (Fp) NotI, 5′– TTGCGGCCGCGGCTCCCTTGCCTGGTCCTCCTGCCGATTC–3′; 5′ homology arm reverse primer (Rp) SalI, 5′–GCGTCGACGGTCGTCTGCTGAGCCCGTGGCCCCGGG–3′; 3′ homology arm Fp ClaI, 5′–GGATCGATTCGAGCTTGCGATTCAGCGGGCACACCTTAGG–3′; and 3′ homology arm Rp AscI, 5′– AAGGCGCGCCAGGGTCTGGCAGCTGAGGAAGTTCCCCTAA–3′. These homology arms were cloned into a vector that incorporated both a neomycin-resistance cassette for positive selection and a diphtheria toxin A (*DT-A*) gene for negative selection. The targeting vector was linearized and electroporated into TT2F ES cells. Recombinant ES clones were isolated after culture in medium containing the G418 antibiotic and screened for proper integration by Southern blotting using the 5′ probe, 3′ probe, and neo cassette sequence. Two clones exhibited proper integration, as validated by genomic sequencing, and were chosen for microinjection into 8-cell stage embryos. The resulting chimeric offspring were crossed to C57BL/6N mice and germ-line transmission was confirmed by Southern blotting and PCR. The floxed PGK-neo cassette was removed by crossing with Meox2-Cre mice (The Jackson Laboratory). Genotyping of *Lin28a* mutant mice was performed by PCR analysis. Genomic DNA was isolated from mouse tail snips. Each tail snip was incubated at 50°C with DirectPCR Lysis reagent with Proteinase K for more than 6 h, followed by heating at 80°C for 1 h, to inactivate Proteinase K. The tail lysate (1 μL) was used as a PCR template. Genotyping PCR was carried out using GoTaq Flexi DNA Polymerase, according to the manufacturer’s protocol. The primer sequences used for *Lin28a* genotyping PCR were as follows: *Lin28a* KO genotyping 1, 5′–TACAAGCCACTGGAACACCA–3′; *Lin28a* KO genotyping 2, 5′–GGGGGTTGGGTCATTGTCTTT–3′; and *Lin28a* KO genotyping 3, 5′– GTTCTGCTGGTAGTGGTCGG–3′.

For CRISPR/Cas9-mediated gene targeting via nonhomologous end joining (Wang et al., 2013, Inui et al., 2014), the guide RNA containing the target sequence of the *Cbx2* CDS (CTGAGCAGCGTGGGCGAGC) was synthesized in vitro using mMESSAGE mMACHINE T7 Kit and was purified using MegaClear Transcription Clean-Up kit, according to the manufacturer’s instructions. A mixture containing 250 ng/μL of guide RNA and hCas9 mRNA was microinjected into the cytoplasm of a 1-cell stage embryo (C57BL/6N background). For genotyping, genomic DNA was isolated from mouse tail snips. Genotyping PCR was carried out using GoTaq Flexi DNA Polymerase, according to the manufacturer’s protocol. The primer sequences used for *Cbx2* genotyping PCR were as follows: *Cbx2* CDS genotyping Fp, 5′–CCCTCTGGCCAAACAATAGCTTTCCGCAGGGACC–3′; and *Cbx2* CDS genotyping Rp, 5′–GCGCCACTTGACCAGGTACTCCAGCTTGCCCTGC–3′. The PCR products were treated with ExoSAP-IT and were then used as a template for direct sequencing. Sequence analysis of the *Cbx2* CDS locus was performed using F0 offspring, and mice that carried frameshift mutations were selected for further analysis.

### HEK293T culture

HEK293T cells were purchased from the American Type Culture Collection (ATCC). Cells were maintained in Dulbecco’s Modified Eagle’s medium (DMEM) supplemented with 10% FBS and antibiotics.

### Establishment of ES-Like Cells

*Lin28a*^−/−^ blastocysts were harvested and cultured on mouse embryonic fibroblasts (MEFs) in ES culture medium (15% FBS, 4.5 g/L of D-glucose, 1× Glutamax, 1 mM sodium pyruvate, 1× nonessential amino acids (NEAAs), 0.1 mM 2-mercaptoethanol, and 1 × 10^4^ units/mL of LIF in DMEM) with 3 μM of CHIR 99021 and 1 μM of PD0325901. Each colony was isolated and expanded, followed by genotyping PCR. Wt and *Lin28a*^−/−^ ES-like cells were stained with NBT/BCIP solution to test for alkaline phosphatase activity. Western blotting and q-PCR analyses were performed for each genotype, as described below.

### Western blotting

Whole-protein extracts from the somites and neural tubes of E9.5 embryos were prepared for western blotting. Samples were separated using 10% SDS–PAGE and blotted onto PVDF membranes. The membranes were first incubated with blocking solution (5% skim milk in TBST) and then incubated with the primary antibody in blocking solution. Membranes were washed in TBST three times for 15 min and incubated with a horseradish peroxidase (HRP)-conjugated secondary antibody in blocking solution. The blots were visualized using Chemi-Lumi One, Immobilon, SuperSignal West Femto Maximum Sensitivity Substrate, and LAS-3000 (Fujifilm), followed by analysis using the Multi Gauge Ver3.2 software. β-actin was measured as an internal control. The antibodies used and their dilutions were listed in Key Resources Table.

### *In situ* hybridization

*Lin28a*^−/−^ embryos and Wt littermates were obtained by intercrossing *Lin28a*^+/–^ mice. Whole-mount *in situ* hybridization was performed as described previously (Yokoyama et al., 2009); the details of the probe sequence can be obtained from the “EMBRYS” website (http://embrys.jp/embrys/html/MainMenu.html). Briefly, embryos were fixed in 4% PFA/PBT and dehydrated in a series of increasing MetOH concentrations.

Rehydrated samples were bleached with 6% H_2_O_2_ in PBT and treated with 10 μg/mL of Protease K for 10 min at room temperature (RT), stopped with 0.2% glycine, and refixed with 4% PFA/0.2% glutaraldehyde in PBT for 20 min at RT. RNA hybridization was performed at 70°C for more than 14 h, after prehybridization for 1 h in hybridization buffer (50% formamide, 5× SSC, 1% SDS, 50 μg/mL of tRNA, and 50 μg/mL of heparin in RNase-free H_2_O). Subsequently, embryos were washed three times in wash buffer 1 (50% formamide, 5× SSC, and 1% SDS in RNase-free H_2_O) and twice in wash buffer 2 (50% formamide, 2× SSC, and 5% Chaps in RNase-free H_2_O). After blocking with 10% sheep serum in TBST for 1 h at RT, samples were incubated with an anti-DIG-AP Fab fragment antibody and 1% sheep serum in TBST overnight (O/N) at 4°C. After a series of washes with TBST, embryos were equilibrated in alkaline phosphatase buffer (NTMT) and developed with NBT/BCIP solution (Roche). After the color reaction, the embryos were rinsed in TBST several times and postfixed in 4% PFA/PBT at 4°C.

*In situ* hybridization of sections was performed on Wt and *Lin28a*^−/−^ embryos at E12.5, as described previously (Uchibe et al., 2012). Embryos were fixed in 4% PFA/PBT, dehydrated in a series of increasing MetOH concentrations, and embedded in paraffin. Sagittal sections (10 μm) were stained with Alcian Blue and Fast Red to outline the pre-vertebrae. Deparaffinized and rehydrated sections were treated with 8 μg/mL of Proteinase K (Roche) in PBS for 10 min, and the reaction was stopped with 0.2% glycine in PBS. After postfixation with 4% PFA, samples were acetylated in acetylation buffer (100 mM triethanolamine, 2.5 mM acetic anhydride; pH was adjusted to 8.0 using HCl). Sections were incubated in prehybridization buffer (50% formamide, 5× SSC) for 1 h at 65°C. Subsequently, hybridization was performed O/N at 65°C using an RNA probe for *Hoxc13* in hybridization buffer (50% formamide, 5× SSC, 10% dextran sulfate, 5× Denhardt’s solution, 0.1 mg/mL of salmon sperm DNA, and 0.25 mg/mL of tRNA). The sections were washed with 0.2× SSC for 3 h at 65°C and rinsed with neutralize tagment (NT) buffer (100 mM Tris-HCl, pH 7.5, 150 mM NaCl) for 5 min. After blocking with 10% sheep serum in NT buffer, samples were incubated with an anti-DIG-AP Fab fragment antibody and 1% sheep serum O/N at 4°C. After a series of washes with NT buffer, samples were equilibrated in NTM (100 mM NaCl, 100 mM Tris-HCl, pH 9.5, and 50 mM MgCl_2_) and developed using an NBT/BCIP solution. After the color reaction, the embryos were counterstained with Fast green.

### Skeletal preparation

Whole-mount skeletal preparations of neonatal mice of each genotype were performed using Alcian Blue and Alizarin Red S staining. For RA treatment, 1 mg/kg of RA was injected intraperitoneally at 7.5 dpc, and the skeletal patterning of each genotype was analyzed at E15.5. The samples were fixed in 100% ethanol (EtOH) for 1–2 days after the majority of the skin and internal organs were removed. The 100% EtOH wash was changed several times. After fixation, the samples were incubated in Alcian Blue solution (0.03% Alcian Blue 8GX, 80% EtOH, and 20% acetic acid) for up to 2 days. The samples were rinsed in distilled water three times and incubated in Alizarin Red Solution (0.01% Alizarin Red S, 1% KOH in H_2_O) O/N. The samples were treated with discoloring solution (1% KOH, 20% glycerol in H_2_O) for 4–7 days. The samples were soaked in a series of glycerol/EtOH solutions (20% glycerol, 20% EtOH; 50% glycerol, 50% EtOH) and stored in 100% glycerol.

### Quantitative PCR

Total RNA was isolated from whole embryos (Figure 3A, 3B), or from dissected somites and neural tubes (Figure 2A, 3C, 3G) at E9.5 using ISOGEN (Nippon Gene), according to the manufacturer’s instructions. For SYBR green q-PCR, a complementary DNA (cDNA) was produced using Superscript II reverse transcriptase, 1 μg of total RNA, and an oligo(dT)18 primer. q-PCR analysis was performed using the SYBR Green PCR Master Mix and an ABI 7900HT instrument (Applied Biosystems). *Gapdh* was measured as an internal control to normalize sample differences. The primer sets used for all *Hox* genes were described by Kondrashov et al. (2011). The primer sequences used for other genes were as follows: *Lin28a* Fp1, 5’–CTCGGTGTCCAACCAGCAGT–3’; *Lin28a* Rp1, 5’–CACGTTGAACCACTTACAGATGC–3’; *Lin28a* Fp2, 5’– AGGCGGTGGAGTTCACCTTTAAGA–3’; *Lin28a* Rp2, 5’– AGCTTGCATTCCTTGGCATGATGG–3’; *Cbx2* Fp, 5’– AGGCCGAGGAAACACACAGT–3’; *Cbx2* Rp, 5’–GGAGGAAGAGGACGAACTGC–3’; *Oct3/4* Fp, 5’–GTTTCTGAAGTGCCCGAAGC–3’; *Oct3/4* Rp, 5’– GCGCCGGTTACAGAACCATA–3’; *Nanog* Fp, 5’–ACCTCAGCCTCCAGCAGATG–3’; *Nanog* Rp, 5’–ACCGCTTGCACTTCATCCTT–3’; *Sox2* Fp, 5’– GGCAGCTACAGCATGATGCAGGAGC–3’; *Sox2* Rp, 5’– CTGGTCATGGAGTTGTACTGCAGG–3’; *Gapdh* Fp, 5’–CCTGGTCACCAGGGCTGC– 3’; and *Gapdh* Rp, 5’–CGCTCCTGGAAGATGGTGATG–3’.

For microRNAs, cDNAs were produced using the TaqMan MicroRNA Reverse Transcription kit according to the manufacturer’s protocol. q-PCR was performed using TaqMan Rodent MicroRNA Array A and B and TaqMan MicroRNA Assays for *let-7a*, *let-7b*, *let-7c*, *let-7d*, *let-7e*, *let-7f*, *let-7g*, *let-7i*, *mir-98*, *mir-10a*, *mir-10b*, *mir-196a*, *mir-196b*, and *RNU6B*. *RNU6B* was measured as an internal control to normalize sample differences.

### Luciferase assay

The pLuc2 reporter vector was as described previously (Miyaki et al., 2010). To create the *let-7* sensor vector, the chemically synthesized *let-7* complementary sequence was annealed and inserted between the EcoRI and XhoI sites. To create the pLuc2-candidate gene 3′UTR vector, the predicted *let-7* target sequence of each genes of 3′UTR was cloned into pLuc2. Fragment containing mutation in *let-7* target sequence were also cloned in pLux2. The miRNA precursor sequence (40 bp) was cloned into pcDNA3.1 and used as an miRNA-expressing vector. Transfection into HEK293T cells was performed using Lipofectamine 2000 or FugeneHD. The transfected cells were incubated for 48 h, and luciferase activity was determined using the Dual-Glo Luciferase Assay System.

### Chromatin immunoprecipitation

Harvested E9.5 embryos were dissected into somites and neural tubes. Genomic DNA was isolated from the yolk sac and genotyping PCR was performed. Samples were cryopreserved until use. For each assay, chromatin immunoprecipitation (ChIP) was performed on a pool of 10 embryos. Each antibody (5 μg) was used for immunoprecipitation. The antibodies used for ChIP were listed in Key Resources Table. The frozen samples were cross-linked with 1% formaldehyde in PBS for 10 min at RT. Cross-linking was stopped by adding 100 μL of 1.25 M glycine for 5 min at RT. Samples were washed with PBS and suspended in cell lysis buffer (10 mM Tris-HCl (pH 7.5), 10 mM NaCl, 3 mM MgCl2, 0.5% NP-40, and 1 mM PMSF). Nuclei were collected by centrifugation and resuspended in cell lysis buffer twice. Samples were suspended in 130 μL of nucleus lysis buffer (50 mM Tris-HCl (pH 8.0), 10 mM EDTA (pH 8.0), 1% SDS, and 1 mM PMSF) and transferred into Covaris microTUBEs. The chromatin was sheared by sonication (peak power, 105; duty factor, 5.0; cycles/burst, 200; duration, 10 min). The sheared DNA was diluted in IP dilution buffer (20 mM Tris-HCl (pH 8.0), 2 mM EDTA (pH 8.0), 150 mM NaCl, 1% Triton X-100, 0.1% SDS, and 1 mM PMSF), added to antibody beads, and rotated O/N at 4°C. Precipitated beads with chromatin were washed four times with ChIP wash buffer 1 (20 mM Tris-HCl (pH 8.0), 2 mM EDTA (pH 8.0), 150 mM NaCl, 1% Triton X-100, 0.1% SDS, and 1 mM PMSF) and twice with ChIP wash buffer 2 (20 mM Tris-HCl (pH 8.0), 2 mM EDTA (pH 8.0), 500 mM NaCl, 1% Triton X-100, 0.1% SDS, and 1 mM PMSF). After washing with TE, chromatin was isolated using nucleus lysis buffer at 65°C. The isolated chromatin was de-cross-linked for 6 h at 65°C. After Proteinase K treatment, DNA was purified using a PCR purification kit (elute in 50 μL of H_2_O). q-PCR was performed on immunoprecipitated DNA and input DNA and analyzed for the efficiency of immunoprecipitation by each antibody. The primer sequences used for ChIP q-PCR were as follows: ChIP *Hoxa1* Fp, 5’– TGAGAAAGTTGGCACGGTCA–3’; ChIP *Hoxa1* Rp, 5’– C ACTGCCAAGGATGGGGTAT–3’; ChIP *Hoxa2* Fp, 5’– CTCCAAGGAGAAGGCCATGA–3’; ChIP *Hoxa2* Rp, 5’– CGACAGGGGGAAAAGATGTC–3’; ChIP *Hoxa3* Fp, 5’– GTTGTCGCTGGAGGTGGAG–3’; ChIP *Hoxa3* Rp, 5’–GCCAGAGGACGCAGGAAAT– 3’; ChIP *Hoxa4* Fp, 5’–AACGACACCGCGAGAAAAAT–3’; ChIP *Hoxa4* Rp, 5’– GGGAACTTGGGCTCGATGTA–3’; ChIP *Hoxa5* Fp, 5’–TCCCCCGAATCCTCTGTATC– 3’; ChIP *Hoxa5* Rp, 5’–ATTGCATTTCCCTCGCAGTT–750 3’; ChIP *Hoxa6* Fp, 5’– GTTCGGCCATCCAGAAACA–3’; ChIP *Hoxa6* Rp, 5’–CCCCTCTGCAGGACTGTGAT– 3’; ChIP *Hoxa7* Fp, 5’–AGCCTTCACCCGACCTATCA–3’; ChIP *Hoxa7* Rp, 5’– AGCACAGCCTCGTTCTCTCC–3’; ChIP *Hoxa9* Fp, 5’– CCTCCCGGGTTAATTTGTAGC–3’; ChIP *Hoxa9* Rp, 5’– CCCCTGCCTTGGTTATCCTT–3’; ChIP *Hoxa10* Fp, 5’– CCTAGACTCCACGCCACCAC–3’; ChIP *Hoxa10* Rp, 5’– GGCTGGAGACAGCTCCTCA–3’; ChIP *Hoxa11* Fp, 5’– AGAGCTCGGCCAACGTCTAC–3’; ChIP *Hoxa11* Rp, 5’– AACTGGTCGAAAGCCTGTGG–3’; ChIP *Hoxa13* Fp, 5’– ACTTCGGCAGCGGCTACTAC–3’; ChIP *Hoxa13* Rp, 5’– CATGTACTTGTCGGCGAAGG–3’; ChIP *Hoxc13* Fp, 5’– CAGGAGACCCAGGCTTAGCA–3’; ChIP *Hoxc13* Rp, 5’– GCATGCGGACACACTTCATT–3’; ChIP *Hoxd12* Fp, 5’– GGAGATGTGTGAGCGCAGTC–3’; ChIP *Hoxd12* Rp, 5’– CTGCCATTGGCTCTCAGGTT–3’.

### Knockdown of *let-7* in ES-like cells

To knockdown *let-7* expression, guide-RNAs targeting the *let-7* family members were constructed. The target sequences of *let-7* family members were as follows: *let-7a-1*, TAGTAGGTTGTATAGTTTT; *let-7a-2* and *let-7c-1*, GGTTGAGGTAGTAGGTTGT; *let-7b*, TAGTAGGTTGTGTGGTTTC; *let-7c-2*, TAGTAGGTTGTATGGTTTT; *let-7d*, TAGTAGGTTGCATAGTTTT; *let-7e*, GTAGGAGGTTGTATAGTTG; *let-7f-1*, TAGTAGATTGTATAGTTGT; *let-7f-2*, TAGTAGATTGTATAGTTTT; *let-7g*, TAGTAGTTTGTACAGTTTG; and *let-7i*, AGGTAGTAGTTTGTGCTGT (see also Figure 5H). Four guide-RNA-expressing plasmid vectors and an hCas9 vector (500 ng each) were transfected into 1 × 10^6^ cells using the Neon transfection system, according to the manufacturer’s instructions. Transfected cells were cultured in ES medium containing 0.5 μg/mL of puromycin for 2 days. Each colony was isolated and expanded, followed by PCR and sequence analysis. The primer sequences used for *let-7* genotyping PCR were as follows: *let-7a-1* Fp, 5′–GGCTTATAGCCCAGGTGTATCAT–3′; *let-7a-1* Rp, 5′– ACTTGCCCATTCCCATCATC–3′; *let-7a-2* Fp, 5′–TTCTTATGAACGGCCCGAGT–3′; *let-7a-2* Rp, 5′–CCGTTGATCACCTGTGTTGC–3′; *let-7c-1* Fp, 5′– TGGTAGGCACAGGCCTTTCT–3′; *let-7c-1* Rp, 5′–CAATGTGTGGTTGGCGATCT–3′; *let-7b* Fp, 5′–TTTGCTCGCTGCTAATGGAA–3′; *let-7b* Rp, 5′– GGCCTCATGGACTCATGACA–3′; *let-7c-2* Fp, 5′–GTCTCCCCGTCTCCCCTTAC–3′; *let-7c-2* Rp, 5′–AGGTGCCCTGAAAATGCTGT–3′; *let-7d* Fp, 5′– TTTGGCTTTTGCCAAGATCA–3′; *let-7d* Rp, 5′–TGCTTTCCAAAACTTCCCAGT–3′; *let-7e* Fp, 5′–TGAATTCCTGGGTTCCTTGG–3′; *let-7e* Rp, 5′– TCAAGATGGCATAGAGACTGCAA–3′; *let-7f-1* Fp, 5′–GATGATGGGAATGGGCAAGT– 3′; *let-7f-1* Rp, 5′–CCAAAAGGCCTGGTCCTAGA–3′; *let-7f-2* Fp, 5′– TCTTGTGTGCTTGTCTCCCATT–3′; *let-7f-2* Rp, 5′–CTGAGAACCACTGCCACCAG– 3′; *let-7g* Fp, 5′–TGGTGTATTTCTTTTGTTGGGTTG–3′; *let-7g* Rp, 5′– TGAACAACTCCAAGCCTCTCA–3′; *let-7i Fp*, 5′–GGGCCCCGGATGTAAGATGG–3′; and *let-7i* Rp, 5′–CCTCGAGAACGAAACCCAAC–3′. The PCR products were treated with ExoSAP-IT (Affimetrix) and used as templates for direct sequencing. Clones of *let-7* family members with deletions of several nucleotides were selected for further analysis. Embryoid bodies were produced from each clone, and expression changes of *Hox* genes were analyzed over 3 days. Cells (1 × 10^6^) were suspended in 1 mL of DMEM with 10% FBS and plated in low-adhesion culture dishes. After several hours, self-aggregated ES-like cells were resuspended in 10 mL of medium. The medium was changed every other day. RNA isolation and q-PCR analysis are described above.

## Statistical analyses

Two-tailed independent Student’s *t*-tests were used to determine all *P* values. Asterisks indicate statistically significant differences (at *p* < 0.05), whereas n.s. indicates an absence of significance.

## Acknowledgements

We thank Dr. Hirohito Shimizu for the technical advice on RA treatment assay, Ms. Moe Tamano for the embryo manipulation, Drs. Satohsi Yamashita and Kazuhiko Nakabayashi for technical advice on ChIP assay, Prof. Mikiko C. Siomi for critical and helpful discussion, and Ms. Izumi A. Tsune and Dr. Spencer J. Spratt for their support in manuscript preparation. We also thank all other Asahara lab members for their support.

## Author contributions

T.S., K.K. and H.A. designed the study, analyzed the data, and wrote the manuscript; T.S., Y.I., Sa.T., and H.U-K. generated the knockout mice; T.S., K.K. and S.Y. analyzed gene expression patterns; T.S., M.I., and Sh.T. performed genome editing; M.M., K.A., and H.A. directed and supervised the study. All authors discussed the data and commented on the manuscript.

## SUPPLEMENTAL FIGURE LEGENDS

**Figure S1.**
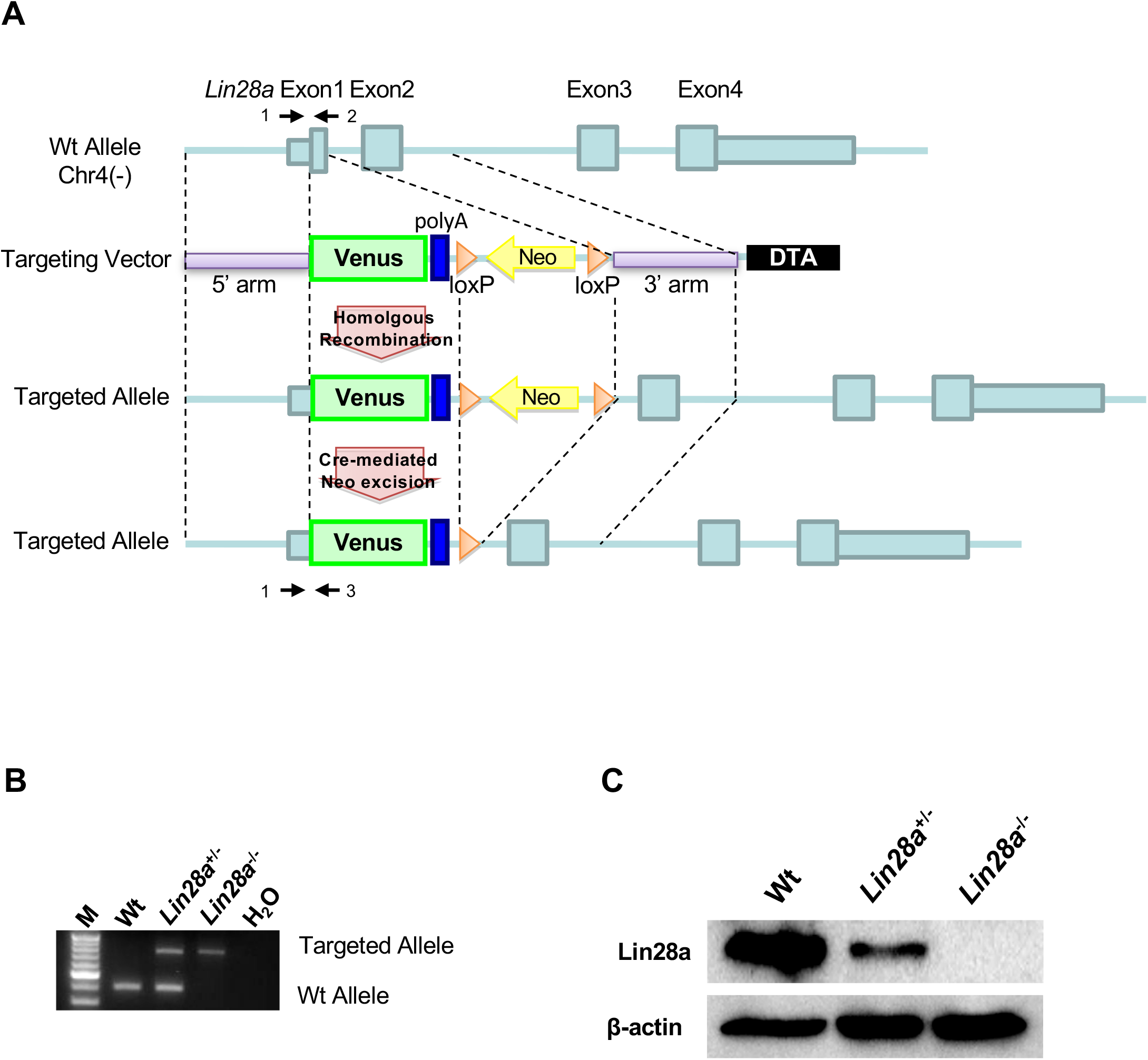
Generation of *Lin28a*^−/−^ mice. (A) Schematic diagram of the Lin28a gene-targeting construct. The small light-blue box indicates the 5′ and 3′ untranslated regions (UTRs), and the large light-blue box indicates the coding region. Arrows (1–3) show the genomic PCR primers used for genotyping. 5′ and 3′ arms, 5′ and 3′ homology arms; Neo, PGK promoter and neomycin-resistance gene; DTA, diphtheria toxin A chain. (B) Genotyping via PCR of Lin28a mutants. The Wt allele produced PCR products of about 400 bp, whereas the targeted allele yielded products of about 750 bp. (C) Western blot analysis of Lin28a in E9.5 whole embryos. β-actin is shown as a loading control.

**Figure S2.**
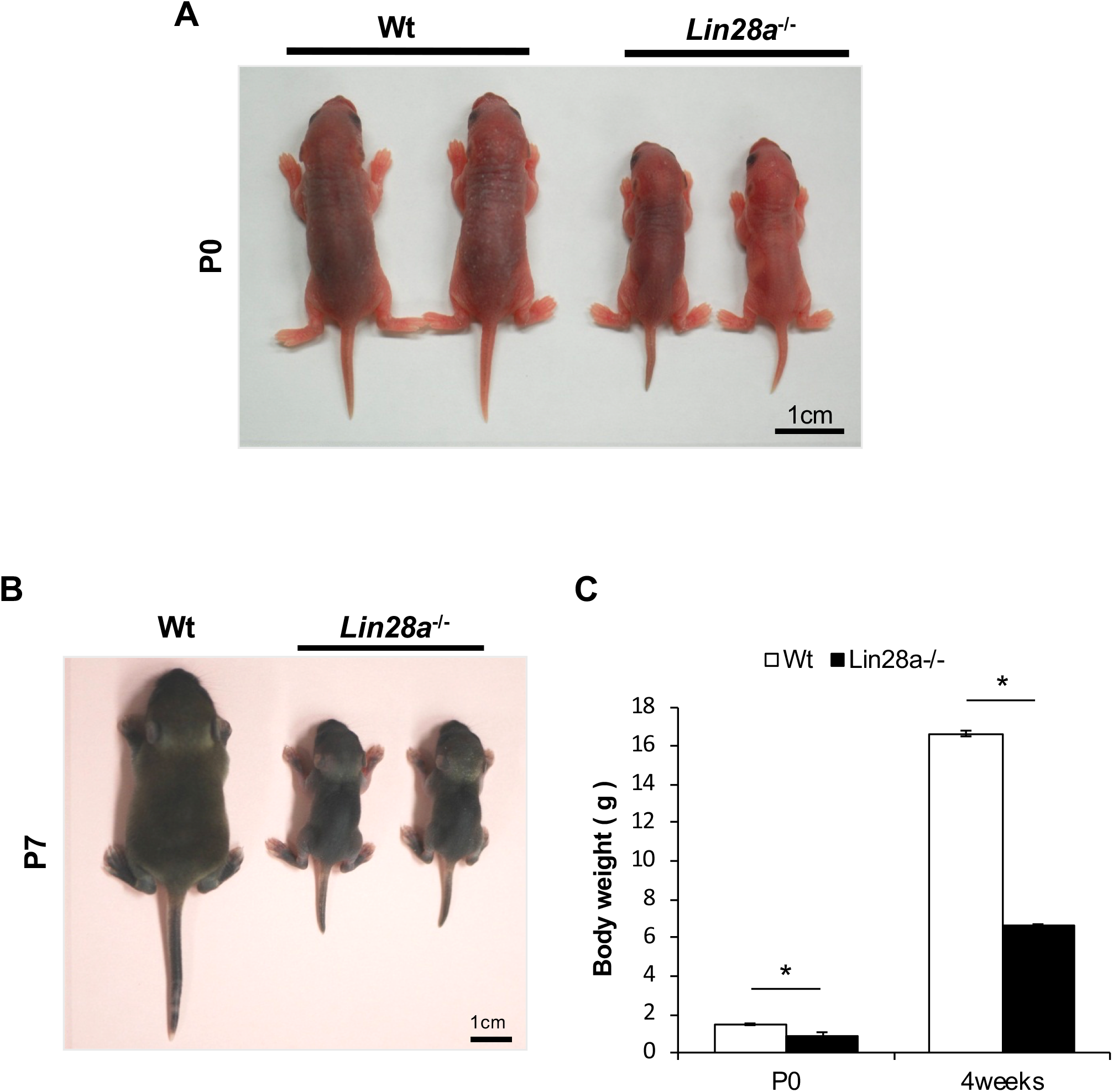
*Lin28a*^−/−^ mice exhibit growth defects. (A, B) Appearance of Wt and *Lin28a*^−/−^ mice. Representative Wt (left) and *Lin28a*^−/−^ (right) pups at P0 (A) and P7 (B) are shown. (C) Body weight of Wt and *Lin28a*^−/−^ mice. Data are expressed as the mean ± SEM (n = 4). *P < 0.05.

**Figure S3.**
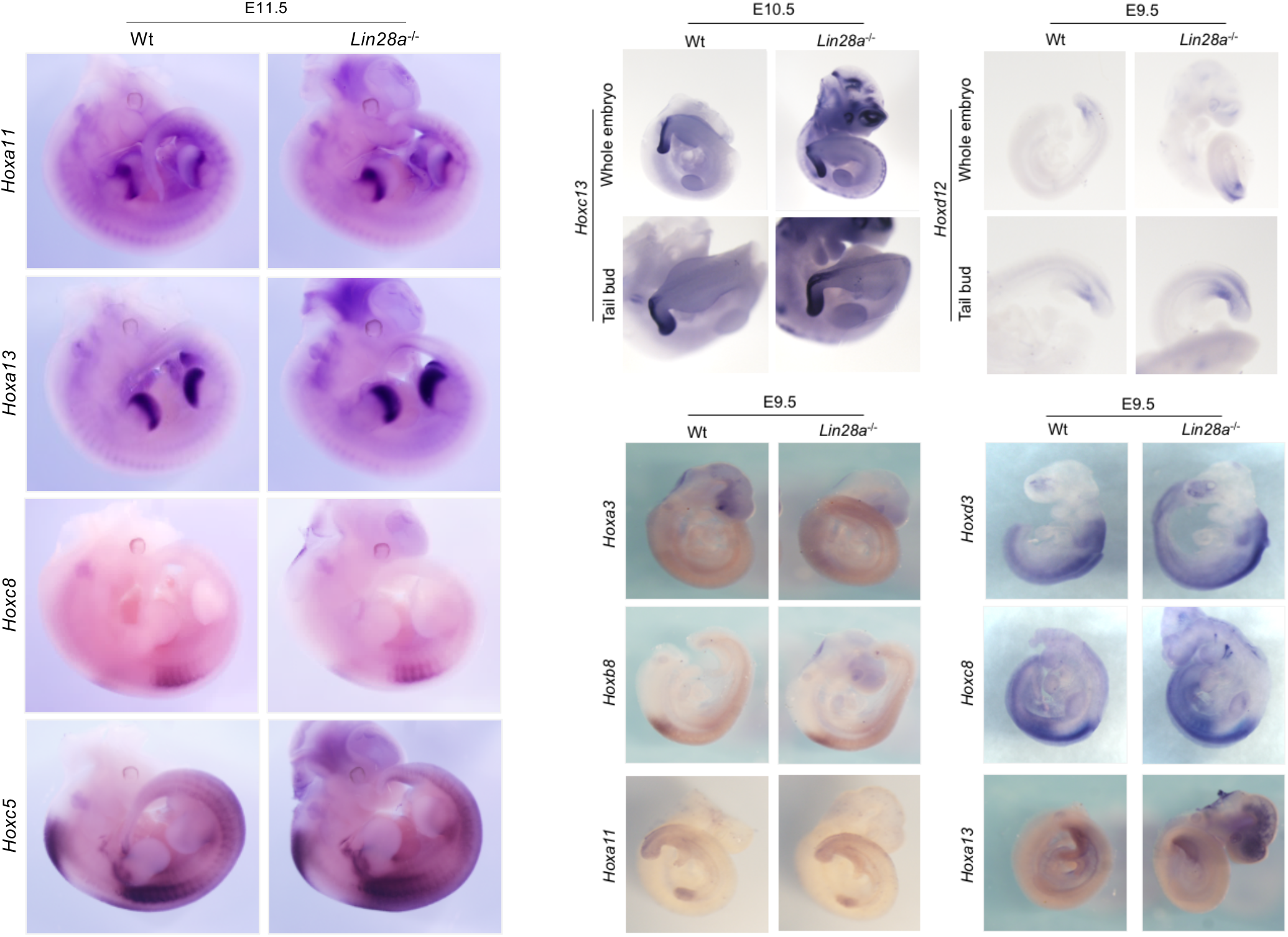
Whole-mount *in situ* hybridization of *Hox* genes in Lin28 knockout embryos (Related to Fig. 2B) Whole-mount *in situ* hybridization of Wt and *Lin28a*^−/−^ mice embryos at E9.5 (using *Hoxa3*, *Hoxd3*, *Hoxb8*, *Hoxc8*, *Hoxa11*, *Hoxd12* and *Hoxa13* probes), at E10.5 (using *Hoxc13* probe) and at E11.5 (using *Hoxc5*, *Hoxc8*, *Hoxa11* and *Hoxa13* probes).

**Figure S4.**
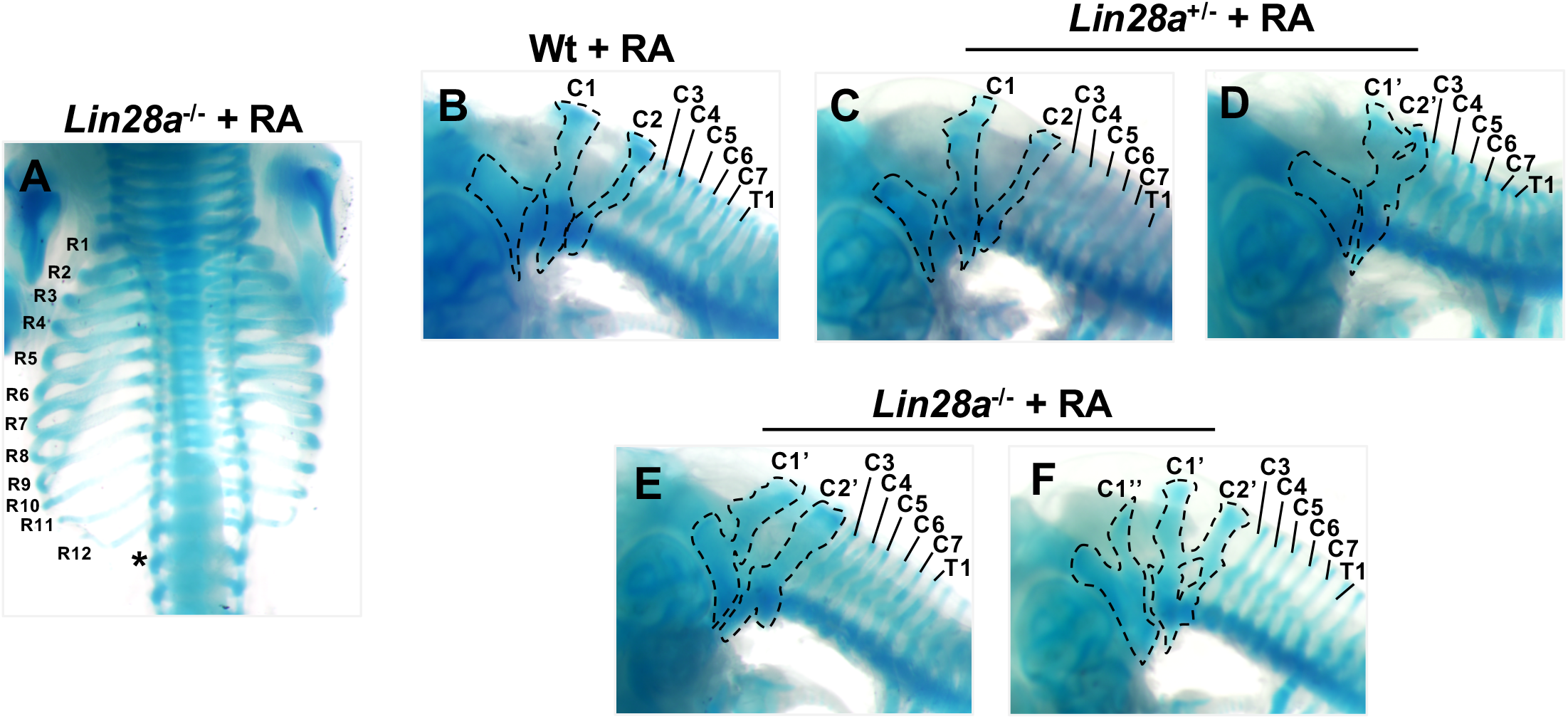
RA sensitivity in *Lin28a* mutant mice. (A) Dorsal views of thoracic vertebrae and ribs of RA-treated *Lin28a*^−/−^ mice. R1–R12, 1^st^ to 12^th^ ribs. The asterisk indicates the ablation of the 13^th^ rib. (B–F) Lateral view of cervical and upper thoracic vertebrae of each genotype treated with RA. C1–C7, 1^st^ to 7^th^ cervical vertebrae; T1, 1^st^ thoracic vertebra. The dotted lines from left to right show the exoccipital bone and C1 and C2, respectively. C1′ and C2′ show fusion and morphological changes of C1 and C2, respectively. C1¢¢ indicates an additional C1 vertebra.

**Figure S5.**
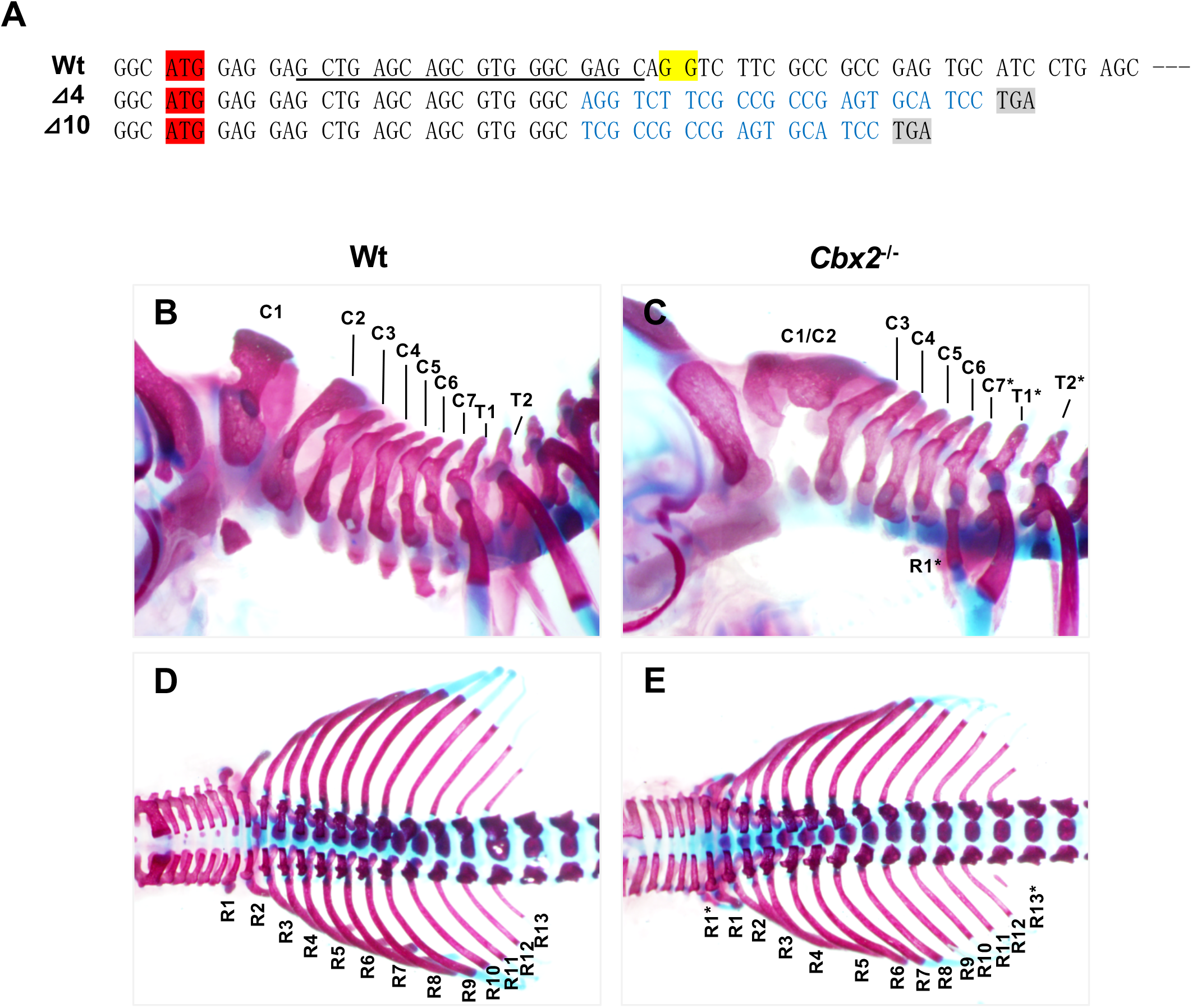
Skeletal defects in *Cbx2* mutant mice. (A) *Cbx2* targeting and the sequence of *Cbx2* mutants. The start codon of Cbx2 is highlighted in red and the PAM sequence for hCas9 is highlighted in yellow. Targeting sequences are underlined. The predicted stop codons in mutants are highlighted in gray. (B–E) Skeletal preparations of Wt and *Cbx2*^−/−^ mice. Lateral views of cervical and upper thoracic vertebrae (B, C) and dorsal views of thoracic vertebrae and ribs (D, E) are shown. C1–C7, 1^st^ to 7^th^ cervical vertebrae; T1 and T2, 1st and 2nd thoracic vertebrae; R1–R13, 1^st^ to 13^th^ ribs; C1/C2, fusion of C1 and C2; the asterisks indicate the posterior transformation of vertebrae.

**Figure S6.**
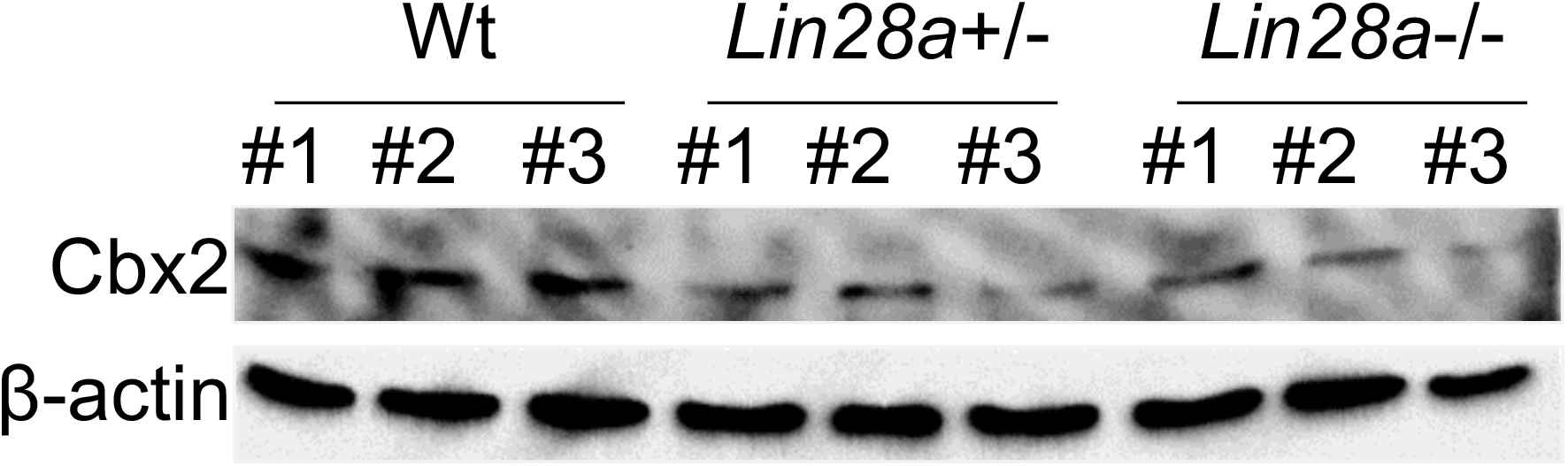
Expression level of Cbx2 in *Lin28a* mutant embryo. Change in the expression of Cbx2 in *Lin28a*^+/–^ and *Lin28a*^−/−^ mice E9.5 embryos were detected by western blot analyses. β-actin is shown as a loading control.

**Supplemental Table 1.**
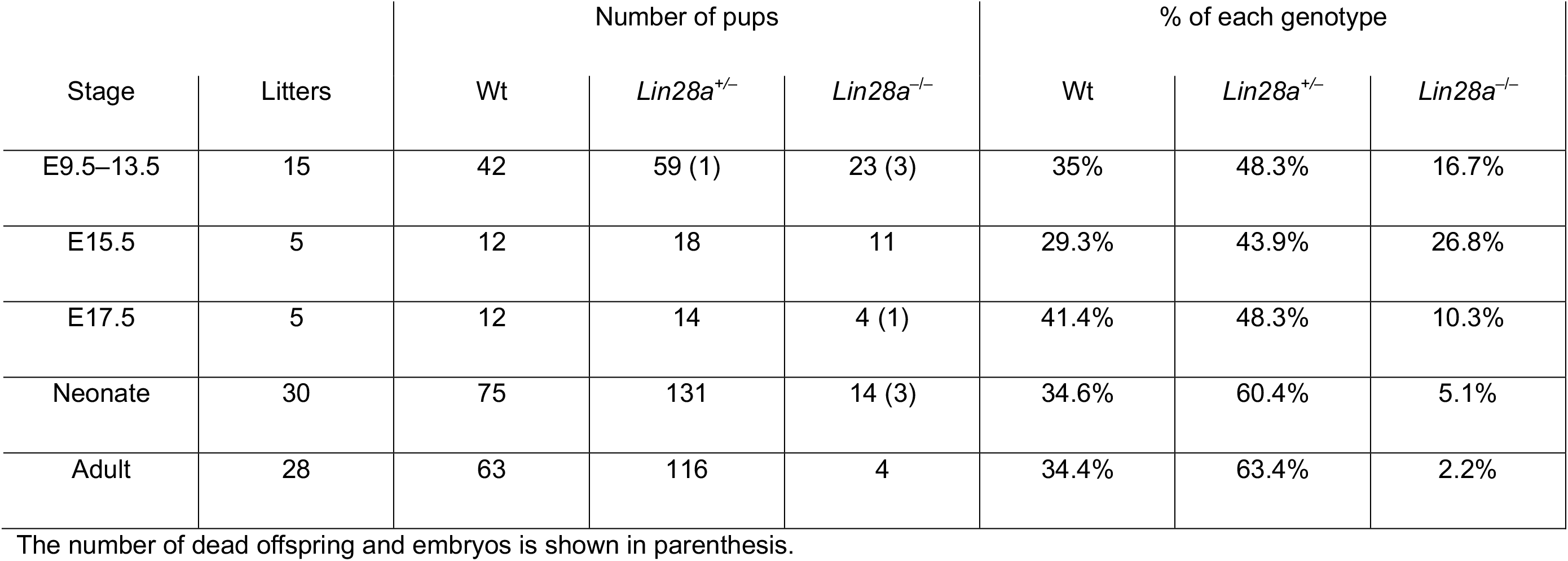
Survival rate of *Lin28a* mutant mice at various stages.

## References

Aires, R., de Lemos, L., Novoa, A., Jurberg, A. D., Mascrez, B., Duboule, D. & Mallo, M.A 2019. Tail Bud Progenitor Activity Relies on a Network Comprising Gdf11, Lin28, and Hox13 Genes. Dev Cell, 48, 383-395.e8.

Akasaka, T., Kanno, M., Balling, R., Mieza, M. A., Taniguchi, M. & Koseki, H. 1996. A role for mel-18, a Polycomb group-related vertebrate gene, during theanteroposterior specification of the axial skeleton. Development, 122, 1513–22.

Ambros, V. & Horvitz, H. 1984. Heterochronic mutants of the nematode Caenorhabditis elegans. Science, 226, 409–416.

Chang, H. M., Triboulet, R., Thornton, J. E. & Gregory, R. I. 2013. A role for the Perlman syndrome exonuclease Dis3l2 in the Lin28-let-7 pathway. Nature, 497, 244–8.

Cho, J., Chang, H., Kwon, S. C., Kim, B., Kim, Y., Choe, J., Ha, M., Kim, Y. K. & Kim, V. N. 2012. LIN28A is a suppressor of ER-associated translation in embryonic stem cells. Cell, 151, 765–777.

Condie, B. G. & Capecchi, M. R. 1994. Mice with targeted disruptions in the paralogous genes hoxa-3 and hoxd-3 reveal synergistic interactions. Nature, 370, 304–7.

Core, N., Bel, S., Gaunt, S. J., Aurrand-Lions, M., Pearce, J., Fisher, A. & Djabali, M. 1997. Altered cellular proliferation and mesoderm patterning in Polycomb-M33-deficient mice. Development, 124, 721–9.

Courel, M., Friesenhahn, L. & Lees, J. A. 2008. E2f6 and Bmi1 cooperate in axial skeletal development. Dev Dyn, 237, 1232–42.

Faas, L., Warrander, F. C., Maguire, R., Ramsbottom, S. A., Quinn, D., Genever, P. & Isaacs, H. V. 2013. Lin28 proteins are required for germ layer specification in Xenopus. Development, 140, 976–86.

Guo, Y., Chen, Y., Ito, H., Watanabe, A., Ge, X., Kodama, T. & Aburatani, H. 2006. Identification and characterization of lin-28 homolog B (LIN28B) in human hepatocellular carcinoma. Gene, 384, 51–61.

Hafner, M., Max, K. E., Bandaru, P., Morozov, P., Gerstberger, S., Brown, M., Molina, H. & Tuschl, T. 2013. Identification of mRNAs bound and regulated by human LIN28 proteins and molecular requirements for RNA recognition. Rna, 19, 613–26.

Han, Y. C., Vidigal, J. A., Mu, P., Yao, E., Singh, I., Gonzalez, A. J., Concepcion, C. P., Bonetti, C., Ogrodowski, P., Carver, B., Selleri, L., Betel, D., Leslie, C. & Ventura, A. 2015. An allelic series of miR-17 approximately 92-mutant mice uncovers functional specialization and cooperation among members of a microRNA polycistron. Nat Genet, 47, 766–75.

Hashimoto, N., Brock, H. W., Nomura, M., Kyba, M., Hodgson, J., Fujita, Y., Takihara, Y., Shimada, K. & Higashinakagawa, T. 1998. RAE28, BMI1, and M33 are members of heterogeneous multimeric mammalian Polycomb group complexes. Biochem Biophys Res Commun, 245, 356-65.

He, C., Kraft, P., Chen, C., Buring, J. E., Pare, G., Hankinson, S. E., Chanock, J., Ridker, P. M., Hunter, D. J. & Chasman, D. I. 2009. Genome-wide association studies identify loci associated with age at menarche and age at natural menopause. Nat Genet, 41, 724–8.

Heimberg, A. & Mcglinn, E. 2012. Building a robust a-p axis. Curr Genomics, 13, 278–88.

Heo, I., Joo, C., Kim, Y. K., Ha, M., Yoon, M. J., Cho, J., Yeom, K. H., Han, J. & Kim, V. N. 2009. TUT4 in concert with Lin28 suppresses microRNA biogenesis through pre-microRNA uridylation. Cell, 138, 696–708.

Hornstein, E., Mansfield, J. H., Yekta, S., Hu, J. K., Harfe, B. D., Mcmanus, M., Baskerville, S., Bartel, D. P. & Tabin, C. J. 2005. The microRNA miR-196 acts upstream of Hoxb8 and Shh in limb development. Nature, 438, 671–4.

Inui, M., Miyado, M., Igarashi, M., Tamano, M., Kubo, A., Yamashita, S., Asahara, H., Fukami, M. & Takada, S. 2014. Rapid generation of mouse models with defined point mutations by the CRISPR/Cas9 system. Sci Rep, 4, 5396.

Johnson, C. D., Esquela-Kerscher, A., Stefani, G., Byrom, M., Kelnar, K., Ovcharenko, D., Wilson, M., Wang, X., Shelton, J., Shingara, J., Chin, L., Brown, D. & Slack, F. J. 2007. The let-7 microRNA represses cell proliferation pathways in human cells. Cancer Res, 67, 7713–22.

Johnson, L., Greenbaum, D., Cichowski, K., Mercer, K., Murphy, E., Schmitt, E., Bronson, R. T., Umanoff, H., Edelmann, W., Kucherlapati, R. & Jacks, T. 1997. K-ras is an essential gene in the mouse with partial functional overlap with N-ras. Genes & Development, 11, 2468–2481.

Johnson, S. M., Grosshans, H., Shingara, J., Byrom, M., Jarvis, R., Cheng, A., Labourier, E., Reinert, K. L., Brown, D. & Slack, F. J. 2005. RAS is regulated by the let-7 microRNA family. Cell, 120, 635–47.

Juan, A. H. & Ruddle, F. H. 2003. Enhancer timing of Hox gene expression: deletion of the endogenous Hoxc8 early enhancer. Development, 130, 4823–34.

Katoh-Fukui, Y., Tsuchiya, R., Shiroishi, T., Nakahara, Y., Hashimoto, N., Noguchi, K. & Higashinakagawa, T. 1998. Male-to-female sex reversal in M33 mutant mice. Nature, 393, 688–92.

Kessel, M. & Gruss, P. 1991. Homeotic transformations of murine vertebrae and concomitant alteration of Hox codes induced by retinoic acid. Cell, 67, 89–104.

Koera, K., Nakamura, K., Nakao, K., Miyoshi, J., Toyoshima, K., Hatta, T., Otani, H., Aiba, A. & Katsuki, M. 1997. K-ras is essential for the development of the mouse embryo. Oncogene, 15, 1151–9.

Kong, D., Heath, E., Chen, W., Cher, M. L., Powell, I., Heilbrun, L., Li, Y., Ali Sethi, S., Hassan, O., Hwang, C., Gupta, N., Chitale, D., Sakr, W. A., Menon, M. & Sarkar, F. H. 2012. Loss of Let-7 Up-Regulates EZH2 in Prostate Cancer Consistent with the Acquisition of Cancer Stem Cell Signatures That Are Attenuated by BR-DIM. PLoS One, 7, e33729.

Lee, Y. S. & Dutta, A. 2007. The tumor suppressor microRNA let-7 represses the HMGA2 oncogene. Genes Dev, 21, 1025–30.

Lettre, G., Jackson, A. U., Gieger, C., Schumacher, F. R., Berndt, S. I., Sanna, S., Eyheramendy, S., Voight, B. F., Butler, J. L., Guiducci, C., Illig, T., Hackett, R., Heid, I. M., Jacobs, K. B., Lyssenko, V., Uda, M., Boehnke, M., Chanock, S. J., Groop, L. C., Hu, F. B., Isomaa, B., Kraft, P., Peltonen, L., Salomaa, V., Schlessinger, D., Hunter, D. J., Hayes, R. B., Abecasis, G. R., Wichmann, H. E., Mohlke, K. L. & Hirschhorn, J. N. 2008. Identification of ten loci associated with height highlights new biological pathways in human growth. Nat Genet, 40, 584–91.

Li, X., Isono, K., Yamada, D., Endo, T. A., Endoh, M., Shinga, J., Mizutani-Koseki, Y., Otte, A. P., Casanova, M., Kitamura, H., Kamijo Sharif, J., Ohara, O., Toyada, T., Bernstein, B. E., Brockdorff, N. & Koseki, H. 2011. Mammalian polycomb-like Pcl2/Mtf2 is a novel regulatory component of PRC2 that can differentially modulate polycomb activity both at the Hox gene cluster and at Cdkn2a genes. Mol Cell Biol, 31, 351–64.

Li, Z., Wang, L., Xu, J. & Yang, Z. 2015. MiRNA expression profile and miRNA-mRNA integrated analysis (MMIA) during podocyte differentiation. Mol Genet Genomics, 290, 863–75.

Ma, W., Ma, J., Xu, J., Qiao, C., Branscum, A., Cardenas, A., Baron, A. T., Schwartz, P., Maihle, N. J. & Huang, Y. 2013. Lin28 regulates BMP4 and functions with Oct4 to affect ovarian tumor microenvironment. Cell Cycle, 12, 88–97.

Madison, B. B., Liu, Q., Zhong, X., Hahn, C. M., Lin, N., Emmett, M. J., Stanger, B. Z., Lee, J. S. & Rustgi, A. K. 2013. LIN28B promotes growth and tumorigenesis of the intestinal epithelium via Let-7. Genes Dev, 27, 2233–45.

Mallo, M. & Alonso, C. R. 2013. The regulation of Hox gene expression during animal development. Development, 140, 3951–63.

Mallo, M., Wellik, D. M. & Deschamps, J. 2010. Hox genes and regional patterning of the vertebrate body plan. Dev Biol, 344, 7–15.

Mayr, C., Hemann, M. T. & Bartel, D. P. 2007. Disrupting the pairing between let-7 and Hmga2 enhances oncogenic transformation. Science, 315, 1576–9.

Morey, L., Pascual, G., Cozzuto, L., Roma, G., Wutz, A., Benitah, S. A. & di Croce, L. 2012. Nonoverlapping functions of the Polycomb group Cbx family of proteins in embryonic stem cells. Cell Stem Cell, 10, 47–62.

Moss, E. G., Lee, R. C. & Ambros, V. 1997. The cold shock domain protein LIN-28 controls developmental timing in C. elegans and is regulated by the lin-4 RNA. Cell, 88, 637–46.

Moss, E. G. & Tang, L. 2003. Conservation of the heterochronic regulator Lin-28, its developmental expression and microRNA complementary sites. Developmental Biology, 258, 432–442.

Nielsen, A. L., Oulad-Abdelghani, M., Ortiz, J. A., Remboutsika, E., Chambon, P. & Losson, R. 2001. Heterochromatin formation in mammalian cells: interaction between histones and HP1 proteins. Mol Cell, 7, 729–39.

O’carroll, D., Erhardt, S., Pagani, M., Barton, S. C., Surani, M. A. & Jenuwein, T. 2001. The polycomb-group gene Ezh2 is required for early mouse development. Mol Cell Biol, 21, 4330–6.

Ong, K. K., Elks, C. E., Li, S., Zhao, J. H., Luan, J., Andersen, L. B., Bingham, S. A., Brage, S., Smith, G. D., Ekelund, U., Gillson, C. J., Glaser, B., Golding, J., Hardy, R., Khaw, K. T., Kuh, D., Luben, R., Marcus, M., Mcgeehin, M. A., Ness, A. R., Northstone, K., Ring, S. M., Rubin, C., Sims, M. A., Song, K., Strachan, D. P., Vollenweider, P., Waeber, G., Waterworth, D. M., Wong, A., Deloukas, P., Barroso, I., Mooser, V., Loos, R. J. & Wareham, N. J. 2009. Genetic variation in LIN28B is associated with the timing of puberty. Nat Genet, 41, 729–33.

Pasquinelli, A. E., Reinhart, B. J., Slack, F., Martindale, M. Q., Kuroda, M. I., Maller, B., Hayward, D. C., Ball, E. E., Degnan, B., Muller, P., Spring, J., Srinivasan, A., Fishman, M., Finnerty, J., Corbo, J., Levine, M., Leahy, P., Davidson, E. & Ruvkun, G. 2000. Conservation of the sequence and temporal expression of let-7 heterochronic regulatory RNA. Nature, 408, 86–9.

Perry, J. R., Stolk, L., Franceschini, N., Lunetta, K. L., Zhai, G., Mcardle, P. F., Smith, A. V., Aspelund, T., Bandinelli, S., Boerwinkle, E., Cherkas, L., Eiriksdottir, G., Estrada, K., Ferrucci, L., Folsom, A. R., Garcia, M., Gudnason, V., Hofman, A., Karasik, D., Kiel, D. P., Launer, L. J., van Meurs, J., Nalls, M. A., Rivadeneira, F., Shuldiner, A. R., Singleton, A., Soranzo, N., Tanaka, T., Visser, J. A., Weedon, M. N., Wilson, S. G., Zhuang, V., Streeten, E. A., Harris, T. B., Murray, A., Spector, T. D., Demerath, E. W., Uitterlinden, A. G. & Murabito, J. M. 2009. Meta-analysis of genome-wide association data identifies two loci influencing age at menarche. Nat Genet, 41, 648–50.

Qiu, C., Ma, Y., Wang, J., Peng, S. & Huang, Y. 2010. Lin28-mediated post-transcriptional regulation of Oct4 expression in human embryonic stem cells. Nucleic Acids Res, 38, 1240–8.

Reinhart, B. J., Slack, F. J., Basson, M., Pasquinelli, A. E., Bettinger, J. C., Rougvie, A. E., Horvitz, H. R. & Ruvkun, G. 2000. The 21-nucleotide let-7 RNA regulates developmental timing in Caenorhabditis elegans. Nature, 403, 901–6.

Robinton, D. A., Chal, J., Lummertz Da Rocha, E., Han, A., Yermalovich, A. V., Oginuma, M., Schlaeger, T. M., Sousa, P., Rodriguez, A., Urbach, A., Pourquie, O. & Daley, G. Q. 2019. The Lin28/let-7 Pathway Regulates the Mammalian Caudal Body Axis Elongation Program. Dev Cell, 48, 396–405.e3.

Rybak, A., Fuchs, H., Smirnova, L., Brandt, C., Pohl, E. E., Nitsch, R. & Wulczyn, F. G. 2008. A feedback loop comprising lin-28 and let-7 controls pre-let-7 maturation during neural stem-cell commitment. Nat Cell Biol, 10, 987–93.

Sampson, V. B., Rong, N. H., Han, J., Yang, Q., Aris, V., Soteropoulos, P., Petrelli, N. J., Dunn, S. P. & Krueger, L. J. 2007. MicroRNA let-7a down-regulates MYC and reverts MYC-induced growth in Burkitt lymphoma cells. Cancer Res, 67, 9762–70.

Shinoda, G., de Soysa, T. Y., Seligson, M. T., Yabuuchi, A., Fujiwara, Y., Huang, P. Y., Hagan, J. P., Gregory, R. I., Moss, E. G. & Daley, G. Q. 2013. Lin28a regulates germ cell pool size and fertility. Stem Cells, 31, 1001–9.

Shyh-Chang, N. & Daley, G. Q. 2013. Lin28: primal regulator of growth and metabolism in stem cells. Cell Stem Cell, 12, 395–406.

Small, K. M. & Potter, S. S. 1993. Homeotic transformations and limb defects in Hox A11 mutant mice. Genes Dev, 7, 2318–28.

Soshnikova, N. 2014. Hox genes regulation in vertebrates. Dev Dyn, 243, 49–58.

Soshnikova, N. & Duboule, D. 2009. Epigenetic temporal control of mouse Hox genes in vivo. Science, 324, 1320–3.

Sulem, P., Gudbjartsson, D. F., Rafnar, T., Holm, H., Olafsdottir, E. J., Olafsdottir, G. H., Jonsson, T., Alexandersen, P., Feenstra, B., Boyd, H. A., Aben, K. K., Verbeek, A. L., Roeleveld, N., Jonasdottir, A., Styrkarsdottir, U., Steinthorsdottir, V., Karason, A., Stacey, S. N., Gudmundsson, J., Jakobsdottir, M., Thorleifsson, G., Hardarson, G., Gulcher, J., Kong, A., Kiemeney, L. A., Melbye, M., Christiansen, C., Tryggvadottir, L., Thorsteinsdottir, U. & Stefansson, K. 2009. Genome-wide association study identifies sequence variants on 6q21 associated with age at menarche. Nat Genet, 41, 734–8.

Suzuki, M., Mizutani-Koseki, Y., Fujimura, Y., Miyagishima, H., Kaneko, T., Takada, Y., Akasaka, T., Tanzawa, H., Takihara, Y., Nakano, M., Masumoto, H., Vidal, M., Isono, K. & Koseki, H. 2002. Involvement of the Polycomb-group gene Ring1B in the specification of the anterior-posterior axis in mice. Development, 129, 4171–83.

Trumpp, A., Refaeli, Y., Oskarsson, T., Gasser, S., Murphy, M., Martin, G. R. & Bishop, J. M. 2001. c-Myc regulates mammalian body size by controlling cell number but not cell size. Nature, 414, 768–73.

Uchibe, K., Shimizu, H., Yokoyama, S., Kuboki, T. & Asahara, H. 2012. Identification of novel transcription-regulating genes expressed during murine molar development. Dev Dyn, 241, 1217–26.

van der Lugt, N. M., Domen, J., Linders, K., van Roon, M., Robanus-Maandag, E., Te Riele, H., van der Valk, M., Deschamps, J., Sofroniew, M., van Lohuizen, M. &, et al. 1994. Posterior transformation, neurological abnormalities, and severe hematopoietic defects in mice with a targeted deletion of the bmi-1 proto-oncogene. Genes Dev, 8, 757–69.

Vinagre, T., Moncaut, N., Carapuco, M., Novoa, A., Bom, J. & Mallo, M. 2010. Evidence for a myotomal Hox/Myf cascade governing nonautonomous control of rib specification within global vertebral domains. Dev Cell, 18, 655–61.

Viswanathan, S. R., Daley, G. Q. & Gregory, R. I. 2008. Selective blockade of microRNA processing by Lin28. Science, 320, 97–100.

Wang, H., Yang, H., Shivalila, C. S., Dawlaty, M. M., Cheng, A. W., Zhang, F. & Jaenisch, R. 2013. One-step generation of mice carrying mutations in multiple genes by CRISPR/Cas-mediated genome engineering. Cell, 153, 910–8.

Wellik, D. M. 2007. Hox patterning of the vertebrate axial skeleton. Dev Dyn, 236, 2454–63.

West, J. A., Viswanathan, S. R., Yabuuchi, A., Cunniff, K., Takeuchi, A., Park, I. H., Sero, J. E., Zhu, H., Perez-Atayde, A., Frazier, A. L., Surani, M. A. & Daley, G. Q. 2009. A role for Lin28 in primordial germ-cell development and germ-cell malignancy. Nature, 460, 909–13.

Widen, E., Ripatti, S., Cousminer, D. L., Surakka, I., Lappalainen, T., Jarvelin, M. R., Eriksson, J. G., Raitakari, O., Salomaa, V., Sovio, U., Hartikainen, A. L., Pouta, A., Mccarthy, M. I., Osmond, C., Kajantie, E., Lehtimaki, T., Viikari, J., Kahonen, M., Tyler-Smith, C., Freimer, N., Hirschhorn, J. N., Peltonen, L. & Palotie, A. 2010. Distinct variants at LIN28B influence growth in height from birth to adulthood. Am J Hum Genet, 86, 773–82.

Wilbert, M. L., Huelga, S. C., Kapeli, K., Stark, T. J., Liang, T. Y., Chen, S. X., Yan, B. Y., Nathanson, J. L., Hutt, K. R., Lovci, M. T., Kazan, H., Vu, A. Q., Massirer, K. B., Morris, Q., Hoon, S. & Yeo, G. W. 2012. LIN28 binds messenger RNAs at GGAGA motifs and regulates splicing factor abundance. Mol Cell, 48, 195–206.

Woltering, J. M. & Durston, A. J. 2008. MiR-10 represses HoxB1a and HoxB3a in zebrafish. PLoS One, 3, e1396.

Xu, B. & Huang, Y. 2009. Histone H2a mRNA interacts with Lin28 and contains a Lin28-dependent posttranscriptional regulatory element. Nucleic Acids Res, 37, 4256–63.

Xu, B., Zhang, K. & Huang, Y. 2009. Lin28 modulates cell growth and associates with a subset of cell cycle regulator mRNAs in mouse embryonic stem cells. Rna, 15, 357–61.

Yang, D. H. & Moss, E. G. 2003. Temporally regulated expression of Lin-28 in diverse tissues of the developing mouse. Gene Expr Patterns, 3, 719–26.

Yang, M., Yang, S. L., Herrlinger, S., Liang, C., Dzieciatkowska, M., Hansen, K. C., Desai, R., Nagy, A., Niswander, L., Moss, E. G. & Chen, J. F. 2015. Lin28 promotes the proliferative capacity of neural progenitor cells in brain development. Development, 142, 1616–1627.

Yekta, S., Shih, I. H. & Bartel, D. P. 2004. MicroRNA-directed cleavage of HOXB8 mRNA. Science, 304, 594–6.

Yokoyama, S., Hashimoto, M., Shimizu, H., Ueno-Kudoh, H., Uchibe, K., Kimura, I. & Asahara, H. 2008. Dynamic gene expression of Lin-28 during embryonic development in mouse and chicken. Gene Expr Patterns, 8, 155–60.

Yokoyama, S., Ito, Y., Ueno-Kudoh, H., Shimizu, H., Uchibe, K., Albini, S., Mitsuoka, K., Miyaki, S., Kiso, M., Nagai, A., Hikata, T., Osada, T., Fukuda, N., Yamashita, S., Harada, D., Mezzano, V., Kasai, M., Puri, P. L., Hayashizaki, Y., Okado, H., Hashimoto, M. & Asahara, H. 2009. A systems approach reveals that the myogenesis genome network is regulated by the transcriptional repressor RP58. Dev Cell, 17, 836–48.

Young, T., Rowland, J. E., van de Ven, C., Bialecka, M., Novoa, A., Carapuco, M., van Nes, J., de Graaff, W., Duluc, I., Freund, J. N., Beck, F., Mallo, M. & Deschamps, J. 2009. Cdx and Hox genes differentially regulate posterior axial growth in mammalian embryos. Dev Cell, 17, 516–26.

Zhang, Z., O’rourke, J. R., Mcmanus, M. T., Lewandoski, M., Harfe, B. D. & Sun, X. 2011. The microRNA-processing enzyme Dicer is dispensable for somite segmentation but essential for limb bud positioning. Dev Biol, 351, 254–65.

Zhou, X., Benson, K. F., Ashar, H. R. & Chada, K. 1995. Mutation responsible for the mouse pygmy phenotype in the developmentally regulated factor HMGI-C. Nature, 376, 771–4.

Zhu, H., Shyh-Chang, N., Segre, A. V., Shinoda, G., Shah, S. P., Einhorn, W. S., Takeuchi, A., Engreitz, J. M., Hagan, J. P., Kharas, M. G., Urbach, A., Thornton, J. E., Triboulet, R., Gregory, R. I., Consortium, D., Investigators, M., Altshuler, D. & Daley, G. Q. 2011. The Lin28/let-7 axis regulates glucose metabolism. Cell, 147, 81–94.

